# Inverting the model of genomics data sharing with the NHGRI Genomic Data Science Analysis, Visualization, and Informatics Lab-space (AnVIL)

**DOI:** 10.1101/2021.04.22.436044

**Authors:** Michael C. Schatz, Anthony A. Philippakis, Enis Afgan, Eric Banks, Vincent J. Carey, Robert J. Carroll, Alessandro Culotti, Kyle Ellrott, Jeremy Goecks, Robert L. Grossman, Ira M. Hall, Kasper D. Hansen, Jonathan Lawson, Jeffrey T. Leek, Anne O’Donnell Luria, Stephen Mosher, Martin Morgan, Anton Nekrutenko, Brian D. O’Connor, Kevin Osborn, Benedict Paten, Candace Patterson, Frederick J. Tan, Casey Overby Taylor, Jennifer Vessio, Levi Waldron, Ting Wang, Kristin Wuichet, AnVIL Team

**Author notes:** AnVIL Team members and affiliations listed on Table 1.

## Abstract

The traditional model of genomic data analysis - downloading data from centralized warehouses for analysis with local computing resources - is increasingly unsustainable. Not only are transfers slow and cost prohibitive, but this approach also leads to redundant and siloed compute infrastructure that makes it difficult to ensure security and compliance of protected data. The NHGRI Genomic Data Science Analysis, Visualization, and Informatics Lab-space (AnVIL; https://anvilproject.org) inverts this model, providing a unified cloud computing environment for data storage, management, and analysis. AnVIL eliminates the need for data movement, allows for active threat detection and monitoring, and provides scalable, shared computing resources that can be acquired by researchers as needed. This presents many new opportunities for collaboration and data sharing that will ultimately lead to scientific discoveries at scales not previously possible.

## I. History of genomics data sharing

Genomics has become a central component to the study of many facets of biology and medicine (Green et al., 2020; Koboldt et al., 2013). Across ancestry analysis (Byrska-Bishop et al., 2021; Karczewski et al., 2020), disease & trait associations (Taliun et al., 2021; Wainschtein et al., 2019), developmental biology (Tanay & Regev, 2017; Trapnell et al., 2014), and many other fields, large scale genome and genomics sequencing has grown tremendously over the past few decades, driven in large part by the technological improvements that have substantially decreased the cost and time required for sequencing (Goodwin et al., 2016). For example, the National Human Genome Research Institute (NHGRI) Centers for Common Disease Genomics (CCDG) and Centers for Mendelian Genomics (CMG) programs seek to identify the genetic components of many major common and rare diseases through the sequencing of more than one hundred thousand genomes (Green et al., 2020). This scale of analysis opens many new opportunities for discovery that would not otherwise be possible, especially for detecting weak associations with rare variants that can only be measured over large cohorts (McCarthy et al., 2008). However, this scale of sequencing also introduces major new technical challenges that require overhauling how genomics and genomics data science are performed. Most urgently, it has become increasingly impractical to perform genomics research by replicating project data across institutional computing clusters, causing us to reformulate how genomics data can be shared and analyzed.

Because the power of genomics is often only realized through large scale data aggregation, genomics has developed a strong tradition for collaborative research and the open sharing of data. Most famously, this tenet was codified by the leaders of the Human Genome Project in 1996 as the “Bermuda Principles”, where they agreed that all human genomic sequence information generated by the project should be made freely available and in the public domain within 24 hours after generation (Barranco, 2021). These principles were established to maximize the benefit of the data to society, especially as private companies during this era were beginning to apply for patents around human gene sequences (Gold & Carbone, 2010). These core principles were later extended in 2009 by the “Toronto Agreement”, which established the rules for sharing data prepublication (Toronto International Data Release Workshop Authors et al., 2009), and later in 2015 by the NIH Genomic Data Sharing Policy, which requires all large scale sequencing data funded by the NIH to be openly shared (National Institutes of Health, 2014). Complementing the efforts by the funding agencies, many major scientific journals now require data to be deposited into public databases before papers will be published, especially journals serving the genomics community (Powell, 2021).

In response to these requirements for data sharing, several large repositories have been established for storing and sharing genomics data. For high throughput sequencing data, the NCBI Sequence Read Archive (SRA) has emerged as the largest publicly available repository, with over 50 Petabases (Pbp) of data currently available through multiple cloud providers and NCBI servers (Leinonen et al., 2011). As most patients have not consented for open release of their genomics data, the closely related database of Genotypes and Phenotypes (dbGaP) was developed to archive and distribute genomics and related data from studies that have investigated the interaction of genotype and phenotype in humans (Tryka et al., 2014). This database currently manages access for 7,582 datasets in 1,232 studies, most of which are “controlled access” where researchers must apply for access to the datasets to an NIH Data Access Committee (DAC) that evaluates if the research goals are consistent with patient consent forms and any constraints identified by the institutions that submitted the data.

However, as valuable as these and related databases have become, they are generally static resources that do not allow detailed analysis to be performed directly within these systems. Instead, researchers using public data and researchers involved in large sequencing efforts most often begin by downloading the data to an institutional computing cluster for analysis.

## II. Inverting the model of data sharing

The traditional model of genomic analysis has been centered around institutional computing clusters where researchers install and maintain their own suite of computational tools to analyze the datasets that are stored directly within their data center. This model presents a high level of flexibility and control for an individual researcher, but the siloed nature of this model introduces several major barriers and inefficiencies. To start, this model leads to redundant infrastructure where each institution establishes its own data center, and creates major administrative inefficiencies where many of the same analysis tools must be deployed and maintained within each center. Software management tools like bioconda (Gruning et al., 2018) or integrated analysis suites such as Bioconductor (Gentleman et al., 2004), aim to simplify such installations, but maintaining software remains a huge burden in aggregate considering the large number of data centers and users involved.

This model is particularly challenging for collaborative analysis, as it requires data to be copied from one data center to another, which becomes more difficult and costly as the data sets increase in size. For example, a moderately large project, such as the 1000 Genomes project, which contains the CRAM files for 3,202 genomes in the extended collection (Byrska-Bishop et al., 2021) is ~80TB and requires several days to make a single copy over typical institutional Internet connections. Larger studies, such as the recent TopMed release with 53,831 genomes (Taliun et al., 2021) is approximately 2PB in size for the CRAM files and will require several weeks to several months to make a single copy. Equally important, reproducibility is very challenging in such a distributed analysis as it becomes increasingly difficult to record the provenance of how files are created across systems. In extreme cases, incompatible or conflicting versions of a tool or dataset could be used by different groups, leading to scientifically invalid results.

A much more scalable model for collaborative research is to invert the model of data sharing: instead of moving data to each researcher, researchers remotely move to the data through the use of cloud computing resources (Langmead & Nellore, 2018; Schatz et al., 2010) (**Figure 1**). This way only a single copy of the data needs to be maintained, which can then be accessed and analyzed by any number of researchers. This model introduces substantial advantages, including reduced redundancy and costs for data storage and greater flexibility in computing resources. Notably, computing in the cloud is “elastic”, meaning that additional computational resources can be dynamically added to match the needs for the analysis to be performed at a given time. Crucially, these resources can also be scaled down when they are not needed after an analysis is complete to limit the costs involved. This model is also much more efficient to manage, as software only needs to be installed or updated in one location for all users to benefit. Finally, centralized services, especially intrusion detection and auditing, can be far more detailed to ensure data security for protected data sets.

**Figure 1.**
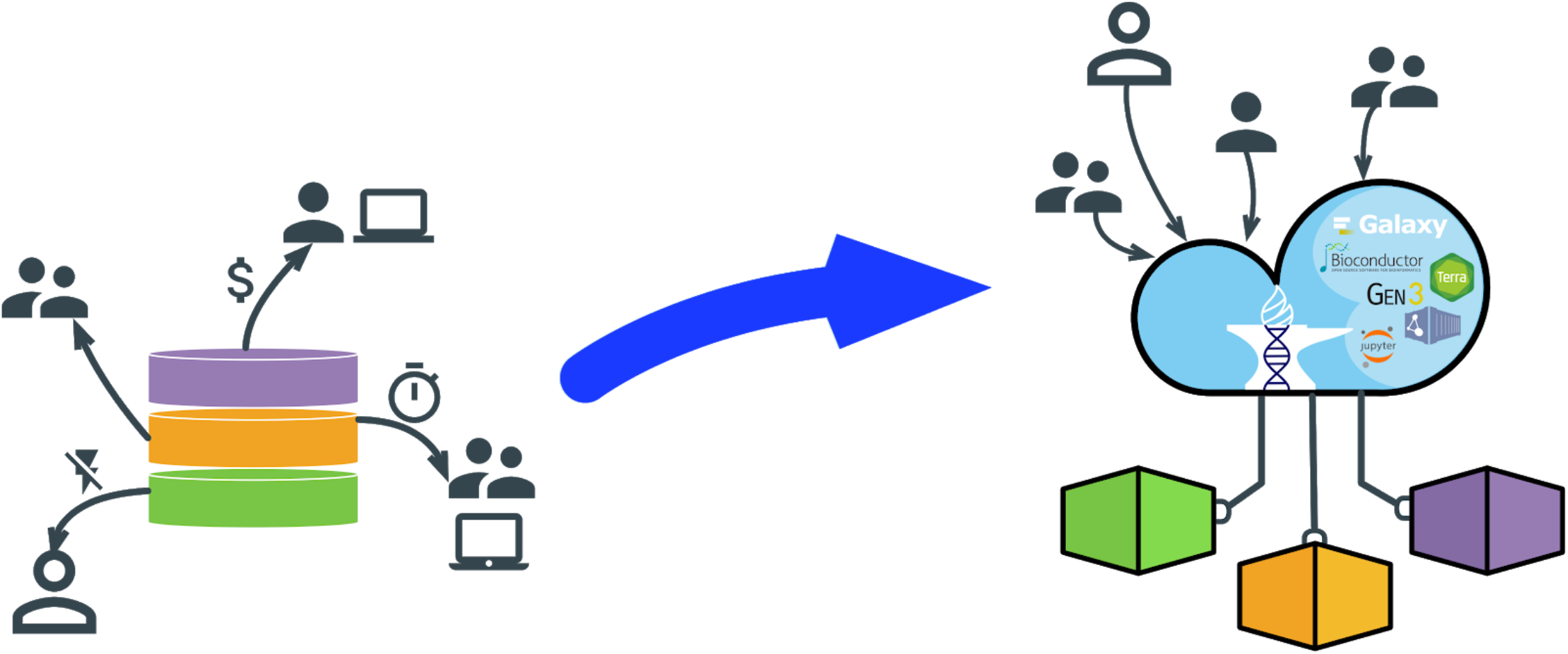
Inverting the model for data sharing. In the traditional model (left), project data are copied from to multiple sites where they are accessed by users on institutional computing clusters. In the inverted model (right), users connect to a cloud-enabled resource such as the AnVIL to remotely access and analyze the data without copying.

Such web-based and cloud-based resources have a strong and growing role in genomics, starting with ubiquitous and classic examples such as the NCBI BLAST server (Johnson et al., 2008) or the UCSC Genome Browser (Navarro Gonzalez et al., 2021). Another rich example is Galaxy (Goecks et al., 2010; Jalili et al., 2020), an open, web-based computational workbench for performing accessible, reproducible, and transparent genomic science with features for executing scientific workflows, data integration, and data and analysis persistence. Even more recent was the NCI Cloud Pilots program, which aimed to provide secure on-demand access to cancer datasets, analysis tools and computing resources (Lau et al., 2017). However, historically these systems have not been sufficient for complete end-to-end analyses of large-scale human genetics and genomics research: their analysis & data management capabilities were too narrow, the data footprint was too large to effectively manage, or the computing environment was not certified for the analysis of protected datasets. Fortunately, as described below, recent advances now make it possible to address all of these issues and provide a cloud-based computing environment that is flexible enough and scalable enough to support any analysis as well as or better than a local computing cluster.

## III. AnVIL System Architecture

In response to these needs, we the AnVIL team with the support of the NHGRI have developed the Genomic Data Science Analysis, Visualization, and Informatics Lab-space (AnVIL). The AnVIL is a federated cloud platform designed to manage and store genomics and related data, enable population-scale analysis, and facilitate collaboration through the sharing of data, code, and analysis results. It includes a variety of graphical user interfaces along with RESTful interfaces and APIs for programmatic access in several popular languages. The compute environment for AnVIL is currently built on the Google Cloud Platform (GCP) to enable massive scalability and capacity for users, as well as a robustly established security perimeter authorized for the storage and analysis of controlled access datasets. Specifically, the AnVIL is a FISMA-Moderate certified computing environment and complies with all requirements set forth in NIST-800-53. This includes robust logging of access to data, periodic audits by third party analysts, and monitoring for abnormal use patterns.

Within the AnVIL, users have several options for analysis and a rich data management ecosystem allowing researchers to search across large collections of data and build novel synthetic cohorts to empower new discoveries out of existing datasets. The analysis components can be broadly characterized as those supporting batch computing, especially through the use of the Workflow Description Language, and interactive computing, using popular analysis suites such as R/Bioconductor, Jupyter notebooks, and Galaxy. Through these components, thousands of genomics analysis tools and workflows are available for a wide variety of analyses. This includes population-scale variant calling with GATK or freebayes (Garrison & Marth, 2012; Van der Auwera et al., 2013), gene expression analysis for both bulk and single cell datasets (Amezquita et al., 2020; Grabherr et al., 2011; Li et al., 2020), methylation analysis (Krueger & Andrews, 2011), COVID-19 viral genomics analysis workflows (Baker et al., 2020; Lemieux et al., 2021), and thousands more. For example, **Supplemental Note 1** displays the workspace for germline variant calling using GATK4. With it, a standard 30x coverage short read dataset can be aligned and variant called in less than 1 day and for less than $5.00 worth of compute. Interestingly, because of the highly scalable nature of cloud computing, processing additional samples, even hundreds or thousands of additional samples, will require approximately the same amount of wall clock time, although costs will scale approximately linearly with the number of samples. **Supplemental Note 2** displays the workspace for analyzing differential gene expression with Bioconductor’s edgeR package. Using the interactive notebooks environment, code and visualizations can easily be interleaved throughout the analysis, starting with quality control through the identification of statistically significant differentially expressed genes. In this example, 1,933 genes decreased expression and 1,794 genes increased expression within a BACH1 knockout dataset. A major current focus is to integrate additional tools and workflows for clinical genomics, especially the calculation and utilization of polygenic risk scores (Torkamani et al., 2018) and pharmacogenomics analysis (Lauschke & Ingelman-Sundberg, 2020). Already more than 15,000 users have used the AnVIL, and the number of users is rapidly growing. Here we review some of the major analysis components found within the AnVIL (**Figure 2**).

**Figure 2:**
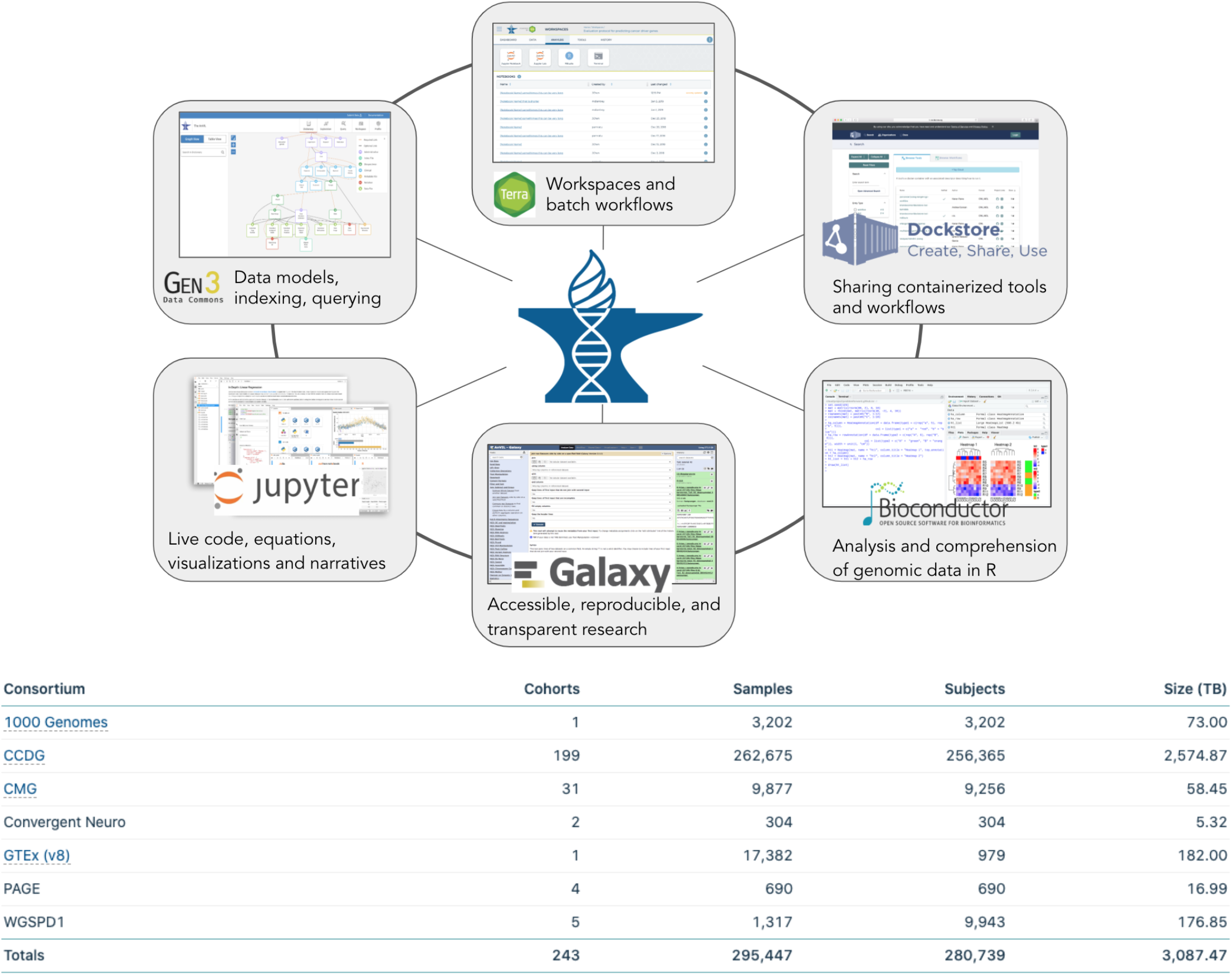
Overview of AnVIL. (top) The AnVIL is a federated cloud environment for the analysis of large genomic and related datasets. The AnVIL is built on a set of established components that have been used in a number of flagship scientific projects. The Terra platform provides a compute environment with secure data and analysis sharing capabilities. Dockstore provides standards based sharing of containerized tools and workflows. R/Bioconductor, Jupyter, and Galaxy provide environments for users at different skill levels to construct and execute analyses. The Gen3 data commons framework provides data and metadata ingest, querying, and organization. (bottom) Summary of the datasets available within the AnVIL as of March 2021 as shown at https://anvilproject.org/data.

### AnVIL Portal: Entry into the AnVIL ecosystem

The initial entry point for AnVIL users is through the AnVIL Portal (https://anvilproject.org). The portal provides unified entry to all of the available tools and datasets available within the system. In addition to a wide variety of training materials and announcements, the portal also has a searchable catalog of the data that are loaded within the AnVIL. Currently, the AnVIL hosts data from >280,000 human genomes from >240 different cohorts spanning CCDG, CMG, the Electronic Medical Records and Genomics (eMERGE) Network, GTEx (GTEx Consortium, 2020) and several others (**Figure 2**). Because only summary information is displayed, any user can browse all of the datasets that are present even if they are not authorized to view the specific data files. This way a user can learn what is available (e.g. all studies of a particular disease or phenotype), and if necessary, direct them to apply for authorization through the appropriate Data Access Committee (e.g. dbGaP or the consortium that maintains the data). The AnVIL also maintains a few critical open access datasets, especially the widely used 1000 Genomes collection of diverse human samples (Byrska-Bishop et al., 2021), including both raw data and harmonized variant calls.

### Gen3: Management, analysis, harmonization, and sharing of large datasets

Gen3 (https://gen3.theanvil.io) is an open-source cloud-based data platform for managing, analyzing, harmonizing, and sharing large datasets. It is based on a set of standards-based services with open APIs called *framework services* for authentication, authorization, creating and accessing FAIR data objects (Wilkinson et al., 2016), and importing and exporting bulk clinical and phenotype data. In particular, it supports assigning persistent digital identifiers to data objects, assigning associated metadata, and accessing the data objects using the GA4GH DRS standard. Gen3 supports authentication and authorization management using OpenID tokens and interoperates with the NIH Research and Authorization Service (RAS). Framework services are also used by other large scale genomics platforms, including NCI’s Cancer Research Data Commons, NHLBI’s BioData Catalyst, and the Kids First Data Resource. Framework services provide the basic scaffolding so that systems such as AnVIL can access data from other cloud-based platforms for genomic data and, in turn, make their data available to these platforms, assuming the appropriate policies supporting this interoperability are in place.

Gen3 also provides services for managing clinical and phenotype data and metadata using a graph database. Gen3’s Windmill service is an interactive website built over the graph database that allows users to explore, submit, and download data. Notably, the Windmill service allows for interactive data exploration, search and cohort-building based on phenotypic variables and data types. Selected cohorts can then be exported into a Terra workspace for downstream processing.

### Terra: Access data, Run analysis tools, and Collaborate in Workspaces

Terra (https://anvil.terra.bio) is a cloud-native platform for biomedical researchers to access data, run analysis tools, and collaborate within the AnVIL. Workspaces are the building blocks of Terra - a dedicated space where collaborators can access and organize the same data and tools and run analyses together. Each workspace is associated with a cloud bucket where data can be stored, such as data generated by a workflow analysis (Reiter et al., 2020) or notebook files for interactive computing. Workspaces also provide data tables for storing and maintaining structured data similar to a spreadsheet. By including links to the data’s actual location in the cloud, the data table links large scale data sets to workspace tools. Finally, within a workspace, users can launch batch analysis jobs or one of several interactive computing environments.

Batch analysis in Terra uses the Workflow Description Language (WDL, https://openwdl.org). WDL is a specialized programming language to specify data processing workflows with a human-readable and -writable syntax. WDL makes it straightforward to define analysis tasks, chain them together in workflows, and parallelize their execution without retooling the application to fit a new programming paradigm. The language makes common patterns (scatter/gather, etc) simple to express, while also admitting uncommon or complicated behavior through conditionals; and strives to achieve portability not only across execution platforms, but also different types of users. WDLs can be stored, shared, and described in Dockstore (described below), and executed in Terra using the Cromwell compute engine (https://cromwell.readthedocs.io) allowing for reproducible analysis of even the largest cohorts with tens of thousands of samples.

### Dockstore: Registry of Tools and Workflows

The Dockstore (https://dockstore.org) provides a place where users can find, share, and use curated tools and workflows. Workflow content is encapsulated in Docker (Boettiger, 2015) and described using a workflow language. The use of Docker makes workflows in Dockstore reproducible by making them easy to run without user installation. Dockstore enables scientists to share analytical tools in a way that makes them machine readable and runnable in a variety of environments. Dockstore currently supports 4 workflow languages: the Workflow Description Language (WDL), Common Workflow Language (CWL), Nextflow, and Galaxy Workflows (GW). Dockstore currently contains 536 workflows in WDL that can be launched in Terra within a few clicks of a button. As such, Dockstore provides one of the most straightforward entry points for users to add batch workflows to AnVIL as it can work with any tool/workflow that can be encapsulated into a Docker container and executed on the command line.

### Jupyter Notebooks: Transparent Code, Visualizations, and Narratives

Jupyter Notebooks (https://jupyter.org) are widely-used open-source web applications that allow users to create and share documents that contain live code, equations, visualizations, and narrative text. Uses include: data cleaning and transformation, numerical simulation, statistical modeling, data visualization, machine learning, and many other analyses. Jupyter supports multiple programming languages, including Python, R, Julia, and Scala. Jupyter Notebooks are an open document format based on JSON that contain a complete record of the user’s sessions and include code, narrative text, equations and rich output. The familiar programming environment makes it easy for new users to perform custom analysis of AnVIL data in a secure and collaborative research environment.

### RStudio: Interactive Machine Learning, Statistical Computing, and Visualizations

RStudio (https://rstudio.com) is an integrated development environment for R, a programming language for statistical computing and visualization. R and its libraries implement a wide variety of statistical and graphical techniques, including linear and nonlinear modeling, classical statistical tests, time-series analysis, classification, clustering, and others. R is easily extensible through functions and extensions, and the R community actively contributes many new packages. Other strengths of R include advanced static and interactive graphics, and facile creation of graphical user interfaces for easy use of highly specialized packages.

### Bioconductor: Community-driven Interactive Genomics with R and RStudio

Bioconductor (https://bioconductor.org) is a free, open source and open development software project for the analysis and comprehension of genomic data, with a focus on developing new computational and statistical methods to interpret biological data. Many of these methods are developed by members of the Bioconductor community (Gentleman et al., 2004), and the Bioconductor project serves as a software repository for a wide range of statistical tools developed in the R programming language. Using a rich array of statistical and graphical features in R, more than 1,900 Bioconductor software packages, 3,200 exemplary experiments, and 50,000 model organism annotation resources have been curated for use in genomic data analysis. The use of these packages requires only an understanding of the R language. As a result, R / Bioconductor packages, which include state-of-the-art statistical inference tools tailored to problems arising in genomics, are widely used by biologists who benefit significantly from their ability to explore and analyze both public and privately developed datasets. Many R / Bioconductor applications can be presented to users in a way that does not require advanced programming expertise, e.g., as ‘Shiny’ applications with graphical interfaces. The AnVIL/Bioconductor environment contains many important resources for the AnVIL, including a fully computable version of the online book Orchestrating Single Cell Analysis with Bioconductor (Amezquita et al., 2020).

### Galaxy: Accessible, Reproducible, and Transparent Genomic Science

Galaxy (http://usegalaxy.org) is an open, web-based computational workbench for performing accessible, reproducible, and transparent genomic science that is used daily by thousands of scientists across the world. There are more than 8,000 analysis tools available within Galaxy that are now accessible within the AnVIL including for variant calling & interpretation, ChIP-seq analysis, RNA-seq analysis, genome assembly, proteomics, epigenomics, transcriptomics, and a host of other analyses in the life sciences. To maintain data security, each AnVIL user runs within an independent Galaxy instance where they can import both unprotected data and the protected human genomics datasets they are authorized to access. This is accomplished using a newly developed import tool allowing data to be added into a user’s instance, where the full suite of Galaxy tools and workflows can be securely run. An AnVIL user can thus use any available Galaxy tool to analyze or visualize data within the boundaries of a compliant, isolated, and secure environment. This marks a major advance as AnVIL users can now leverage Galaxy for the analysis of protected human datasets, which is not possible with other public instances of Galaxy.

### Extending the AnVIL

In addition to the components described above, there are many ways to extend the AnVIL to include new capabilities. The most straightforward approaches are to develop a new Docker-based WDL that can launch novel analysis tools or to wrap an analysis or visualization tool so that it can be executed within the Galaxy GUI. More sophisticated integrations are also possible, using a variety of low level APIs and resources. Recent efforts have focused on deploying new applications using Kubernetes (https://kubernetes.io), which can be used for managing very complicated software stacks on scalable infrastructure. Applications are deployed and managed in the Kubernetes cluster by Helm (https://helm.sh/) in the form of Charts. In this design, a Helm chart translates an application’s software stack into customizable Kubernetes manifests. This model, originally developed by the Galaxy Team to enable Galaxy’s deployment within the AnVIL, can be replicated and extended to facilitate the integration of other platforms of varying complexity into the AnVIL. We also have several major additional components in development, including deploying seqr (https://seqr.broadinstitute.org) and the UCSC Genome Browser (Navarro Gonzalez et al., 2021) within the AnVIL.

## IV. Data Access and Data Use

A key priority of the AnVIL is ensuring responsible data sharing, which includes secure access to the data in its cloud storage and compute environments. The AnVIL Data Access Working Group (DAWG) defines the methods used to securely control and grant access to controlled-access datasets hosted within AnVIL, and is testing improved processes for handling data access requests (DARs). The DAWG evaluates the data coming into AnVIL and considers downstream data access needs. For example, the DAWG generated the Consortium Guidelines for AnVIL Data Access (https://anvilproject.org/learn/for-consortia/consortium-data-access-guidelines) to clarify expectations for the various consortia using AnVIL facilitate inter-consortium data sharing and access controls.

Importantly, the DAWG is leading a pilot of the Data Use Oversight System (DUOS) (https://duos.broadinstitute.org/), a platform developed by the Broad Institute, which aims to expedite data access for researchers, by facilitating and enhancing data access committee’s workflows. The pilot currently includes multiple NIH DACs, who are testing the system and providing feedback to further develop the DUOS software, most notably DUOS’ DAR decision-support algorithm. This algorithm leverages the Global Alliance for Genomics and Health (GA4GH) Data Use Ontology (DUO) (https://github.com/EBISPOT/DUO) to code both datasets’ data use terms and researchers’ proposed research contained within DARs. With both of these inputs in terms from the same ontology, the algorithm can assess if the proposed research is within the bounds of the data use terms, and provide a recommended decision to the DAC. In the long term, the pilot will also provide powerful empirical and conceptual evidence of the feasibility of semi-automated approaches to data use oversight.

The DAWG is also refining the Library Card concept, by which an institution can pre-authorize trusted researchers to make controlled-data access requests. This concept will leverage the GA4GH Passport Visa specification (https://github.com/ga4gh-duri/ga4gh-duri.github.io). If implemented, the Library Card concept would reduce the steps required for researchers to submit a DAR, while ensuring the researcher has the appropriate permissions to do so.

If successful, we believe DUOS and the Library Card concept will standardize and streamline the DAR process. As the number of requests for data increases in magnitude over the years, DUOS could ensure DAC members’ time is reserved for fine-grained judgement of complex requests and the Library Card could streamline the authorization of researchers. We hope that by pioneering implementations of the GA4GH DUO and Passports standards, AnVIL will drive interoperable, ethical, and accelerated genomics research.

## V. AnVIL Community

The AnVIL is designed to support a broad range of user communities, from multi-institution consortia, to individual labs and their trainees, to computational tool developers, and researchers at institutions without access to high-performance computing. Some needs of these communities are common - the ability to upload, manage, and share controlled access protected data, the ability to do high-performance computation in either workflow or interactive environments, and the ability to develop training materials and share results with the broader community. However, the diversity of the AnVIL user base also requires satisfying specific needs of the constituent communities.
- **Consortia and data generators**-The primary needs of these groups include data ingestion, quality control, management and sharing within consortium members and collaborators. We have developed a process for data ingestion and management on the AnVIL platform that supports consortia to share their data while ensuring user management and access via access groups following consortium’s data sharing and access guidelines. As of February 2021, the AnVIL contained over 200 NHGRI datasets including the popular GTEx version 8 data, which is also optionally available for direct download free of egress charges.
- **Research groups and investigators** - The primary needs of these groups include access to data, interactive and batch workflow computing environments, and the ability to manage their data science projects. We have developed a user management system leveraging the Terra workflow and workspace access management system. We have also partnered with STRIDES (https://datascience.nih.gov/strides) to support several pilot user education events with an eye toward scaling support to the broader research community. As of February 2021, the AnVIL has supported computation from more than 1,950 users running more than 775 workflows and launching more than 240 workspaces.
- **Computational tool developers** - tool developers need an environment where they can reproducibly test their genomic data science tools, integrate them into workflows, and share them with the broader community. The AnVIL supports several major avenues of deployment, including Docker containers to execute as WDL workflows, conda packages that can execute within Galaxy, or new Bioconductor packages. Notably, by leveraging existing data science tool developer communities thousands of Bioconductor software packages and Galaxy workflows are already integrated in the AnVIL environment.
- **Under-resourced Genomic Data Science Communities** - one of the biggest advantages of a fully cloud-based computational environment like the AnVIL is the ability to do high-performance computing from anywhere. Genomic data science with the AnVIL is accessible to anyone with a web browser and an internet connection - extending access to high-performance computing to communities that do not have local resources to support this kind of science. We have begun a collaboration called the Genomic Data Science Community Network (http://gdscn.org) with community colleges, historically black colleges and universities, and tribal colleges to support data intensive genomic research and teaching using the AnVIL.

## VI. Interoperability with other cloud platforms

Cloud-based systems such as the NHGRI AnVIL as well as NCI’s CRDC and Cloud Resources and NHLBI’s BioData Catalyst have shifted the way researchers work with genomic and other large omics datasets. Freed from the constraint of needing to download data to local compute infrastructure, these environments have allowed researchers to streamline data access and focus on the analysis to be done. Across the AnVIL and peer projects including NHLBI’s BioData Catalyst (BDCat, https://biodatacatalyst.nhlbi.nih.gov), Common Fund’s Gabriella Miller Kids First Pediatric Research Program (GMFK, https://kidsfirstdrc.org), and NCI Cancer Research Data Commons (CRDC, https://datacommons.cancer.gov) for example, almost eight petabytes of genomic and related data are accessible to researchers in cloud-based analysis platforms and growing quickly. Each of these platforms hosts unique datasets and offers unique analysis components to serve their respective research communities. Yet, despite the enormous opportunity to cross-analyze data from these resources, researchers are faced with a daunting task of understanding the various technical interface differences between systems in order to analyze across them -- from a programmatic, user interface, and even policy perspective. As a result, there is great motivation for these systems to adopt consistent conventions and standards, enabling interoperability that facilitates researchers ability to ask questions across the individual platforms.

The AnVIL project has pushed the interoperability envelope by piloting new technologies and adopting key standards and conventions, from known standards bodies such as the Global Alliance for Genomics and Health (GA4GH, https://www.ga4gh.org). This was done to realize the vision of researchers using data and compute across NIH cloud-based platforms seamlessly. AnVIL’s interoperability strategy focuses on 4 distinct areas: 1) data access, 2) portable analysis, 3) authentication & authorization, and 4) search and handoff between systems. For data access, AnVIL has implemented the GA4GH Data Repository Service (DRS), which provides a consistent interface to data resources on cloud environments (both public and private). To enable portable analysis, AnVIL supports both the Workflow Definition Language (WDL) and Galaxy Workflows through Terra and Galaxy, respectively. Each system allows researchers to write analysis tools and workflows that leverage Docker images, a popular containerization technology that facilitates portability. These workflows are shared through Dockstore which, itself, supports the GA4GH Tool Registry Service (TRS), making it possible to share workflows between many different systems beyond AnVIL. For authentication and authorization the AnVIL uses the Research Auth Service (RAS) from NIH. This uses the OIDC/OAuth2 standards and leverages GA4GH Passports, providing a consistent way to describe datasets a researcher is authorized to access. Finally, AnVIL has explored search and data discovery through the Fast Healthcare Interoperability Resource (FHIR) standard and developed a search handoff mechanism between the AnVIL data discovery portal and Terra analysis environment using the Portal Format for Bioinformatics (PFB).

The interoperability vision and accomplishments of AnVIL were not done in isolation but as part of a larger collaboration within the NIH. The NIH Cloud Platform Interoperability (NCPI, https://anvilproject.org/ncpi) effort was started in late 2019 with the goal of establishing and implementing guidelines and technical standards to empower end-users to analyze data across participating platforms and to facilitate the realization of a trans-NIH, federated data & compute ecosystem spanning AnVIL, BDCat, CRDC, and GMFK, along with strong ties to other NIH services such as dbGaP and the SRA. The NCPI Systems Interoperation working group has focused on leveraging the interoperability standards of DRS and TRS, conventions like PFB, and the auth services of RAS to address real-world scientific use cases. In 2020 these interoperability standards were put to use and researchers were able to search across two of the participating stacks (BDCat and AnVIL), access data from both in the AnVIL workspace environment, parameterize and run a workflow, and generate a meaningful result. The NCPI is using this demonstration as the foundation for future work in 2021 with the goal of expanding the interoperability between systems to all of the NCPI (adding GMKF and CRDC) and beyond.

Beyond technical interoperability, is the need for semantic interoperability. The AnVIL hosts data from a wide diversity of projects which contain differing levels of annotation, alternate ontologies, and even mismatched measurement units, sometimes even within the same project. These issues make analyzing phenotypic data in conjunction with the genomic data difficult, even if the systems utilize the same APIs for data transfer. In developing a unified data transform for the AnVIL data dashboard, we encountered known problems with common metadata and phenotypic data elements, such as inconsistent or missing: project disease mapping, subject disease affected status, consistent keys for specimen, specimen attributes, CRAM/BAM statistics, and other missing files.

Extending on these data normalization efforts researchers from the AnVIL project began mapping common elements to other other NIH projects. One example of NIH investment in these standards is HL7 Fast Healthcare Interoperability Resources (FHIR, http://hl7.org/fhir) - a standard for representing, searching, and sharing clinical data. For data commons, FHIR can be seen as a target data model and staging database for data interchange efforts. With a common data model, protocol and search mechanism for clinical attributes in place, the gap becomes a difference between meta-data values and ontologies. Recent Notices from NHGRI and the NIH at large have emphasized the importance for projects to follow standards for meta-data formatting (https://grants.nih.gov/grants/guide/notice-files/NOT-HG-21-022.html), including the use of FHIR (https://grants.nih.gov/grants/guide/notice-flles/NOT-OD-19-122.html). And while the original primary focus of FHIR development was on enabling the exchange of medical data, its formalization of record versioning, provenance tracking and ontology mapping make it a useful platform for cross project data interoperability. This technology is now being utilized to provide more normalized data models that allow for queries across AnVIL and other NIH projects, such as the Kid’s First Data Resource (https://kidsfirstdrc.org/).

## VII. Outlook

The last twenty years have seen tremendous growth in genomics, with millions of human genomes sequenced so far and many millions more to be sequenced in the near future (Stephens et al., 2015). These data, combined with ever growing amounts of single cell & functional genomics data, electronic medical records, and other healthcare data have the potential to substantially enhance our understanding of the basic processes for healthy life as well as revolutionize the treatment of disease. This research will be accomplished, in part, by aggregating and synthesizing data using new computational, statistical and machine learning methods, combined with new high throughput experimental methods that can systematically evaluate large numbers of candidate relationships. However, reaching these ambitious goals requires us to embrace new paradigms for computational research where cloud computing plays a central role; there is simply no other way to effectively share and analyze data at these scales.

The AnVIL launched just over two years ago. While there has been remarkable progress since then, there is still significant work required before the promise of this effort is fully realized. We are still in the earliest stages of the cloud transformation within the life sciences, and institutions have already made significant investments into institutional computing clusters and data centers that we cannot ignore. During this transition period it is likely that cloud resources will be used for the largest analyses and collaborative research projects, but summarized data and institutionally generated private data will still be analyzed locally. As such, one of the key requirements for the AnVIL is that all of the major analysis components can be run locally: WDLs are increasingly used on institutional computing clusters, R/Bioconductor works equally well on a laptop or in the cloud, and Galaxy can be deployed on a laptop or within an institutional cluster as needed. Within AnVIL we also provide free egress for one of the most important datasets, the raw data for the widely studied GTEx dataset, by mirroring our cloud copy within an academic computing center so that authorized users can access it freely over Internet2 (https://anvilproject.org/news/2020/11/20/nhgri-anvil-now-supports-free-export-of-gtex-data)

Another major consideration for cloud-based research are the costs involved. Even if the cost per genome or cost per sample is only a few dollars, once multiplied by thousands of genomes, the total costs for an analysis can quickly become a major expense. While computing costs also play a major role for local computing, these costs are often amortized or supplemented through institution-wide or department-wide initiatives beyond individual research labs. Researchers currently have limited information for the expected costs for running analysis tools in the cloud, which challenges budgeting and prevents many researchers from adopting cloud solutions. In addition, software developers may not focus on optimizing costs for cloud environments, which increases expense even when relatively simple optimizations are available. We and the entire genomics community must address this using all options available, including (1) educating users of the expected costs for different analyses and strategies for minimizing costs; (2) implementing additional technology safeguards (“bumpers”) that will prevent users from uncontrolled spending; and (3) developing optimized tools and workflows to reduce costs. As cloud costs are broadly a function of CPU time, peak RAM required, and disk space, this will include optimizations for decreasing computing time by leveraging parallel & vectorized computing instructions (e.g. AVX512 vectorization (Darby et al., 2020)) or advanced search strategies (e.g. learned index structures (Kirsche et al., 2020; Kraska et al., 2017)), decreasing RAM requirements using more advanced data structures (e.g. Burrows-Wheeler transform (Langmead et al., 2009), Bloom filters (Chikhi & Rizk, 2013), or Sequence Bloom Trees (Solomon & Kingsford, 2016)), and decreasing storage requirements by using compressed data formats (e.g. CRAM (Hsi-Yang Fritz et al., 2011)), using optimized IO routines (e.g. fixed length records instead of variable length records (Langmead et al., 2019)), and removing intermediate data. This will also include developing heuristics and approximation techniques that can often run substantially faster than more exhaustive approaches (Ondov et al., 2016; Rhyker Ranallo-Benavidez et al., 2020).

Overall, the future of the AnVIL and the future of cloud computing in genomics is very bright. We have several major initiatives underway to enhance our capabilities for basic science and clinical genomics, such as by integrating the tools and data from the Telomere-to-Telomere (Miga et al., 2020) and Human Reference Pan Genome projects to provide more comprehensive and more diverse reference human genomes as well as major efforts with the American Heart Association (AHA), the Electronic Medical Records and Genomics (eMERGE) Network and the Clinical Sequencing Evidence-Generating Research (CSER) programs to increase our capabilities for clinical genomics. We are particularly excited by how these efforts will allow us to consider additional variant types, especially complex structural variants, and additional functional genomics data types, including at the single cell level level, to develop a better understanding of the molecular basis of health and disease across diverse patient populations. Internally, we also have several major technical enhancements planned, such as offering multicloud support, and enhanced support for deploying additional complex applications using kubernetes. In addition, NHGRI is broadening the support for the AnVIL by promoting it as a primary data sharing platform and/or a primary data analysis platform for several funding opportunities. Finally, these efforts are coupled with major efforts for training and outreach to ensure everyone is aware of the platform and can use it for their research needs for many years to come.

**Table 1.**
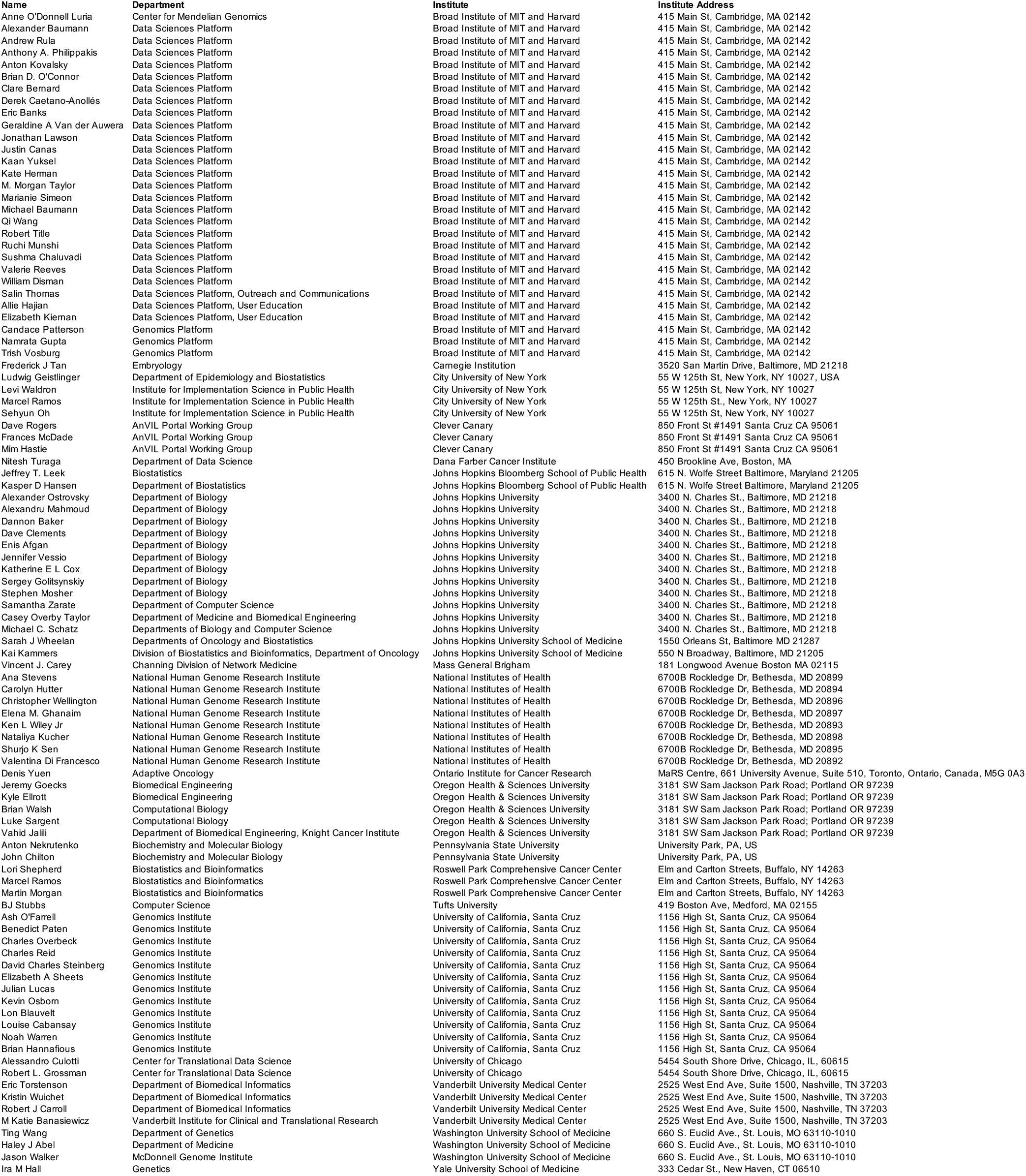
AnVIL Team.

## Acknowledgements

This work is dedicated to the late James Peter Taylor, the Ralph S. O’Connor Professor of Biology and Computer Science at Johns Hopkins University, who was one of the original architects for the AnVIL and an ardent champion for open science (https://galaxyproject.org/jxtx). V.D.F., E.M.G., C.H., N.K., S.K.S., A.S, C.W., and K.L.W. provided substantial involvement and guidance to the project activities and contributed to this manuscript in their official roles as program coordinators for the National Institutes of Health, National Human Genome Research Institute. The AnVIL is supported through cooperative agreement awards from NHGRI with co-funding from OD/ODSS to the Broad Institute (#U24HG010262) and Johns Hopkins University (#U24HG010263). The GDSCN is supported through a contract to Johns Hopkins University (75N92020P00235).

## References

Amezquita, R. A., Lun, A. T. L., Becht, E., Carey, V. J., Carpp, L. N., Geistlinger, L., Marini, F., Rue-Albrecht, K., Risso, D., Soneson, C., Waldron, L., Pages, H., Smith, M. L., Huber, W., Morgan, M., Gottardo, R., & Hicks, S. C. (2020). Orchestrating single-cell analysis with Bioconductor. Nature Methods, 17(2), 137–145. https://doi.org/10.1038/s41592-019-0654-x

Baker, D., van den Beek, M., Blankenberg, D., Bouvier, D., Chilton, J., Coraor, N., Coppens, F., Eguinoa, I., Gladman, S., Gruning, B., Keener, N., Lariviere, D., Lonie, A., Kosakovsky Pond, S., Maier, W., Nekrutenko, A., Taylor, J., & Weaver, S. (2020). No more business as usual: Agile and effective responses to emerging pathogen threats require open data and open analytics. PLoS Pathogens, 16(8), e1008643. https://doi.org/10.1371/journal.ppat.1008643

Barranco, C. (2021). The Human Genome Project. Nature Research. https://doi.org/10.1038/d42859-020-00101-9

Boettiger, C. (2015). An introduction to Docker for reproducible research. ACM SIGOPS Operating Systems Review, 49(1), 71–79. https://doi.org/10.1145/2723872.2723882

Byrska-Bishop, M., Evani, U. S., Zhao, X., Basile, A. O., Abel, H. J., Regier, A. A., Corvelo, A., Clarke, W. E., Musunuri, R., Nagulapalli, K., Fairley, S., Runnels, A., Winterkorn, L., Lowy-Gallego, E., The Human Genome Structural Variation Consortium, Flicek, P., Germer, S., Brand, H., Hall, I. M.,. Zody, M. C. (2021). High coverage whole genome sequencing of the expanded 1000 Genomes Project cohort including 602 trios. In Cold Spring Harbor Laboratory (p. 2021.02.06.430068). https://doi.org/10.1101/2021.02.06.430068

Chikhi, R., & Rizk, G. (2013). Space-efficient and exact de Bruijn graph representation based on a Bloom filter. Algorithms for Molecular Biology: AMB, 8(1), 22. https://doi.org/10.1186/1748-7188-8-22

Darby, C. A., Gaddipati, R., Schatz, M. C., & Langmead, B. (2020). Vargas: heuristic-free alignment for assessing linear and graph read aligners. Bioinformatics. https://doi.org/10.1093/bioinformatics/btaa265

Garrison, E., & Marth, G. (2012). Haplotype-based variant detection from short-read sequencing. In arXiv [q-bio.GNJ. arXiv. http://arxiv.org/abs/1207.3907

Gentleman, R. C., Carey, V. J., Bates, D. M., Bolstad, B., Dettling, M., Dudoit, S., Ellis, B., Gautier, L., Ge, Y., Gentry, J., Hornik, K., Hothorn, T., Huber, W., Iacus, S., Irizarry, R., Leisch, F., Li, C., Maechler, M., Rossini, A. J.,. Zhang, J. (2004). Bioconductor: open software development for computational biology and bioinformatics. Genome Biology, 5(10), R80. https://doi.org/10.1186/gb-2004-5-10-r80

Goecks, J., Nekrutenko, A., Taylor, J., & Galaxy Team. (2010). Galaxy: a comprehensive approach for supporting accessible, reproducible, and transparent computational research in the life sciences. Genome Biology, 11(8), R86. https://doi.org/10.1186/gb-2010-11-8-r86

Gold, E. R., & Carbone, J. (2010). Myriad Genetics: In the eye of the policy storm. Genetics in Medicine: Official Journal of the American College of Medical Genetics, 12(4 Suppl), S39–S70. https://doi.org/10.1097/GIM.0b013e3181d72661

Goodwin, S., McPherson, J. D., & McCombie, W. R. (2016). Coming of age: ten years of next-generation sequencing technologies. Nature Reviews. Genetics, 17(6), 333–351. https://doi.org/10.1038/nrg.2016.49

Grabherr, M. G., Haas, B. J., Yassour, M., Levin, J. Z., Thompson, D. A., Amit, I., Adiconis, X., Fan, L., Raychowdhury, R., Zeng, Q., Chen, Z., Mauceli, E., Hacohen, N., Gnirke, A., Rhind, N., di Palma, F., Birren, B. W., Nusbaum, C., Lindblad-Toh, K.,. Regev, A. (2011). Full-length transcriptome assembly from RNA-Seq data without a reference genome. Nature Biotechnology, 29(7), 644–652. https://doi.org/10.1038/nbt.1883

Green, E. D., Gunter, C., Biesecker, L. G., Di Francesco, V., Easter, C. L., Feingold, E. A., Felsenfeld, A. L., Kaufman, D. J., Ostrander, E. A., Pavan, W. J., Phillippy, A. M., Wise, A. L., Dayal, J. G., Kish, B. J., Mandich, A., Wellington, C. R., Wetterstrand, K. A., Bates, S. A., Leja, D.,. Manolio, T. A. (2020). Strategic vision for improving human health at The Forefront of Genomics. Nature, 586(7831), 683–692. https://doi.org/10.1038/s41586-020-2817-4

Gruning, B., Dale, R., Sjodin, A., Chapman, B. A., Rowe, J., Tomkins-Tinch, C. H., Valieris, R., Koster, J., & Bioconda Team. (2018). Bioconda: sustainable and comprehensive software distribution for the life sciences. Nature Methods, 15(7), 475–476. https://doi.org/10.1038/s41592-018-0046-7

GTEx Consortium. (2020). The GTEx Consortium atlas of genetic regulatory effects across human tissues. Science, 369(6509), 1318–1330. https://doi.org/10.1126/science.aaz1776

Hsi-Yang Fritz, M., Leinonen, R., Cochrane, G., & Birney, E. (2011). Efficient storage of high throughput DNA sequencing data using reference-based compression. Genome Research, 21(5), 734–740. https://doi.org/10.1101/gr.114819.110

Jalili, V., Afgan, E., Gu, Q., Clements, D., Blankenberg, D., Goecks, J., Taylor, J., & Nekrutenko, A. (2020). The Galaxy platform for accessible, reproducible and collaborative biomedical analyses: 2020 update. Nucleic Acids Research, 48(W1), W395–W402. https://doi.org/10.1093/nar/gkaa434

Johnson, M., Zaretskaya, I., Raytselis, Y., Merezhuk, Y., McGinnis, S., & Madden, T. L. (2008). NCBI BLAST: a better web interface. Nucleic Acids Research, 36(Web Server issue), W5–W9. https://doi.org/10.1093/nar/gkn201

Karczewski, K. J., Francioli, L. C., Tiao, G., Cummings, B. B., Alfoldi, J., Wang, Q., Collins, R. L., Laricchia, K. M., Ganna, A., Birnbaum, D. P., Gauthier, L. D., Brand, H., Solomonson, M., Watts, N. A., Rhodes, D., Singer-Berk, M., England, E. M., Seaby, E. G., Kosmicki, J. A.,. MacArthur, D. G. (2020). The mutational constraint spectrum quantified from variation in 141,456 humans. Nature, 581(7809), 434–443. https://doi.org/10.1038/s41586-020-2308-7

Kirsche, M., Das, A., & Schatz, M. C. (2020). Sapling: Accelerating Suffix Array Queries with Learned Data Models. Bioinformatics. https://doi.org/10.1093/bioinformatics/btaa911

Koboldt, D. C., Steinberg, K. M., Larson, D. E., Wilson, R. K., & Mardis, E. R. (2013). The next-generation sequencing revolution and its impact on genomics. Cell, 155(1), 27–38. https://doi.org/10.1016/j.cell.2013.09.006

Kraska, T., Beutel, A., Chi, E. H., Dean, J., & Polyzotis, N. (2017). The Case for Learned Index Structures. In arXiv [cs.DBJ. arXiv. http://arxiv.org/abs/1712.01208

Krueger, F., & Andrews, S. R. (2011). Bismark: a flexible aligner and methylation caller for Bisulfite-Seq applications. Bioinformatics, 27(11), 1571–1572. https://doi.org/10.1093/bioinformatics/btr167

Langmead, B., & Nellore, A. (2018). Cloud computing for genomic data analysis and collaboration. Nature Reviews. Genetics, 19(4), 208–219. https://doi.org/10.1038/nrg.2017.113

Langmead, B., Trapnell, C., Pop, M., & Salzberg, S. L. (2009). Ultrafast and memory-efficient alignment of short DNA sequences to the human genome. Genome Biology, 10(3), R25. https://doi.org/10.1186/gb-2009-10-3-r25

Langmead, B., Wilks, C., Antonescu, V., & Charles, R. (2019). Scaling read aligners to hundreds of threads on general-purpose processors. Bioinformatics, 35(3), 421–432. https://doi.org/10.1093/bioinformatics/bty648

Lau, J. W., Lehnert, E., Sethi, A., Malhotra, R., Kaushik, G., Onder, Z., Groves-Kirkby, N., Mihajlovic, A., DiGiovanna, J., Srdic, M., Bajcic, D., Radenkovic, J., Mladenovic, V., Krstanovic, D., Arsenijevic, V., Klisic, D., Mitrovic, M., Bogicevic, I., Kural, D.,. Seven Bridges CGC Team. (2017). The Cancer Genomics Cloud: Collaborative, Reproducible, and Democratized-A New Paradigm in Large-Scale Computational Research. Cancer Research, 77(21), e3–e6. https://doi.org/10.1158/0008-5472.CAN-17-0387

Lauschke, V. M., & Ingelman-Sundberg, M. (2020). Emerging strategies to bridge the gap between pharmacogenomic research and its clinical implementation. NPJ Genomic Medicine, 5, 9. https://doi.org/10.1038/s41525-020-0119-2

Leinonen, R., Sugawara, H., Shumway, M., & International Nucleotide Sequence Database Collaboration. (2011). The sequence read archive. Nucleic Acids Research, 39(Database issue), D19–D21. https://doi.org/10.1093/nar/gkq1019

Lemieux, J. E., Siddle, K. J., Shaw, B. M., Loreth, C., Schaffner, S. F., Gladden-Young, A., Adams, G., Fink, T., Tomkins-Tinch, C. H., Krasilnikova, L. A., DeRuff, K. C., Rudy, M., Bauer, M. R., Lagerborg, K. A., Normandin, E., Chapman, S. B., Reilly, S. K., Anahtar, M. N., Lin, A. E.,. MacInnis, B. L. (2021). Phylogenetic analysis of SARS-CoV-2 in Boston highlights the impact of superspreading events. Science, 371(6529). https://doi.org/10.1126/science.abe3261

Li, B., Gould, J., Yang, Y., Sarkizova, S., Tabaka, M., Ashenberg, O., Rosen, Y., Slyper, M., Kowalczyk, M. S., Villani, A.-C., Tickle, T., Hacohen, N., Rozenblatt-Rosen, O., & Regev, A. (2020). Cumulus provides cloud-based data analysis for large-scale single-cell and single-nucleus RNA-seq. Nature Methods, 17(8), 793–798. https://doi.org/10.1038/s41592-020-0905-x

McCarthy, M. I., Abecasis, G. R., Cardon, L. R., Goldstein, D. B., Little, J., Ioannidis, J. P. A., & Hirschhorn, J. N. (2008). Genome-wide association studies for complex traits: consensus, uncertainty and challenges. Nature Reviews. Genetics, 9(5), 356–369. https://doi.org/10.1038/nrg2344

Miga, K. H., Koren, S., Rhie, A., Vollger, M. R., Gershman, A., Bzikadze, A., Brooks, S., Howe, E., Porubsky, D., Logsdon, G. A., Schneider, V. A., Potapova, T., Wood, J., Chow, W., Armstrong, J., Fredrickson, J., Pak, E., Tigyi, K., Kremitzki, M.,. Phillippy, A. M. (2020). Telomere-to-telomere assembly of a complete human X chromosome. Nature, 585(7823), 79–84. https://doi.org/10.1038/s41586-020-2547-7

National Institutes of Health. (2014). Final NIH Genomic Data Sharing Policy. In Federal Register (No. 2014-20385; Vol. 79, pp. 51345–51354). https://www.federalregister.gov/d/2014-20385

Navarro Gonzalez, J., Zweig, A. S., Speir, M. L., Schmelter, D., Rosenbloom, K. R., Raney, B. J., Powell, C. C., Nassar, L. R., Maulding, N. D., Lee, C. M., Lee, B. T., Hinrichs, A. S., Fyfe, A. C., Fernandes, J. D., Diekhans, M., Clawson, H., Casper, J., Benet-Pages, A., Barber, G. P.,. Kent, W. J. (2021). The UCSC Genome Browser database: 2021 update. Nucleic Acids Research, 49(D1), D1046–D1057. https://doi.org/10.1093/nar/gkaa1070

Ondov, B. D., Treangen, T. J., Melsted, P., Mallonee, A. B., Bergman, N. H., Koren, S., & Phillippy, A. M. (2016). Mash: fast genome and metagenome distance estimation using MinHash. Genome Biology, 17(1), 132. https://doi.org/10.1186/s13059-016-0997-x

Powell, K. (2021). The broken promise that undermines human genome research. Nature, 590(7845), 198–201. https://doi.org/10.1038/d41586-021-00331-5

Reiter, T., Brooks, P. T., Irber, L., Joslin, S. E. K., Reid, C. M., Scott, C., Titus Brown, C., & Tessa Pierce, N. (2020). Streamlining Data-Intensive Biology With Workflow Systems. In Cold Spring Harbor Laboratory (p. 2020.06.30.178673). https://doi.org/10.1101/2020.06.30.178673

Rhyker Ranallo-Benavidez, T., Lemmon, Z., Soyk, S., Aganezov, S., Salerno, W. J., McCoy, R. C., Lippman, Z. B., Schatz, M. C., & Sedlazeck, F. J. (2020). SVCollector: Optimized sample selection for cost-efficient long-read population sequencing. In Cold Spring Harbor Laboratory (p. 2020.08.06.240390). https://doi.org/10.1101/2020.08.06.240390

Schatz, M. C., Langmead, B., & Salzberg, S. L. (2010). Cloud computing and the DNA data race. Nature Biotechnology, 28(7), 691–693. https://doi.org/10.1038/nbt0710-691

Solomon, B., & Kingsford, C. (2016). Fast search of thousands of short-read sequencing experiments. Nature Biotechnology, 34(3), 300–302. https://doi.org/10.1038/nbt.3442

Stephens, Z. D., Lee, S. Y., Faghri, F., Campbell, R. H., Zhai, C., Efron, M. J., Iyer, R., Schatz, M. C., Sinha, S., & Robinson, G. E. (2015). Big Data: Astronomical or Genomical? PLoS Biology, 13(7), e1002195. https://doi.org/10.1371/journal.pbio.1002195

Taliun, D., Harris, D. N., Kessler, M. D., Carlson, J., Szpiech, Z. A., Torres, R., Taliun, S. A. G., Corvelo, A., Gogarten, S. M., Kang, H. M., Pitsillides, A. N., LeFaive, J., Lee, S.-B., Tian, X., Browning, B. L., Das, S., Emde, A.-K., Clarke, W. E., Loesch, D. P.,. Abecasis, G. R. (2021). Sequencing of 53,831 diverse genomes from the NHLBI TOPMed Program. Nature, 590(7845), 290–299. https://doi.org/10.1038/s41586-021-03205-y

Tanay, A., & Regev, A. (2017). Scaling single-cell genomics from phenomenology to mechanism. Nature, 541(7637), 331–338. https://doi.org/10.1038/nature21350

Torkamani, A., Wineinger, N. E., & Topol, E. J. (2018). The personal and clinical utility of polygenic risk scores. Nature Reviews. Genetics, 19(9), 581–590. https://doi.org/10.1038/s41576-018-0018-x

Toronto International Data Release Workshop Authors, Birney, E., Hudson, T. J., Green, E. D., Gunter, C., Eddy, S., Rogers, J., Harris, J. R., Ehrlich, S. D., Apweiler, R., Austin, C. P., Berglund, L., Bobrow, M., Bountra, C., Brookes, A. J., Cambon-Thomsen, A., Carter, N. P., Chisholm, R. L., Contreras, J. L.,. Yu, J. (2009). Prepublication data sharing. Nature, 461(7261), 168–170. https://doi.org/10.1038/461168a

Trapnell, C., Cacchiarelli, D., Grimsby, J., Pokharel, P., Li, S., Morse, M., Lennon, N. J., Livak, K. J., Mikkelsen, T. S., & Rinn, J. L. (2014). The dynamics and regulators of cell fate decisions are revealed by pseudotemporal ordering of single cells. Nature Biotechnology, 32(4), 381–386. https://doi.org/10.1038/nbt.2859

Tryka, K. A., Hao, L., Sturcke, A., Jin, Y., Wang, Z. Y., Ziyabari, L., Lee, M., Popova, N., Sharopova, N., Kimura, M., & Feolo, M. (2014). NCBI’s Database of Genotypes and Phenotypes: dbGaP. Nucleic Acids Research, 42(Database issue), D975–D979. https://doi.org/10.1093/nar/gkt1211

Van der Auwera, G. A., Carneiro, M. O., Hartl, C., Poplin, R., Del Angel, G., Levy-Moonshine, A., Jordan, T., Shakir, K., Roazen, D., Thibault, J., Banks, E., Garimella, K. V., Altshuler, D., Gabriel, S., & DePristo, M. A. (2013). From FastQ data to high confidence variant calls: the Genome Analysis Toolkit best practices pipeline. Current Protocols in Bioinformatics / Editoral Board, Andreas D. Baxevanis … [et Al.J, 43, 11.10.1–33. https://doi.org/10.1002/0471250953.bi1110s43

Wainschtein, P., Jain, D. P., Yengo, L., Zheng, Z., TOPMed Anthropometry Working Group, Trans-Omics for Precision Medicine Consortium, Adrienne Cupples, L., Shadyab, A. H., McKnight, B., Shoemaker, B. M., Mitchell, B. D., Psaty, B. M., Kooperberg, C., Roden, D., Darbar, D., Arnett, D. K., Regan, E. A., Boerwinkle, E., Rotter, J. I., Allison, M. A.,. Visscher, P. M. (2019). Recovery of trait heritability from whole genome sequence data. In Cold Spring Harbor Laboratory (p. 588020). https://doi.org/10.1101/588020

Wilkinson, M. D., Dumontier, M., Aalbersberg, I. J. J., Appleton, G., Axton, M., Baak, A., Blomberg, N., Boiten, J.-W., da Silva Santos, L. B., Bourne, P. E., Bouwman, J., Brookes, A. J., Clark, T., Crosas, M., Dillo, I., Dumon, O., Edmunds, S., Evelo, C. T., Finkers, R.,. Mons, B. (2016). The FAIR Guiding Principles for scientific data management and stewardship. Scientific Data, 3, 160018. https://doi.org/10.1038/sdata.2016.18

