## Supplemental Notes for "Inverting the model of genomics data sharing with the NHGRI Genomic Data Science Analysis, Visualization, and Informatics Lab-space (AnVIL)"

Supplemental Note 1. GATK4 Germline Variant Calling within Terra

POWERED BY Terra | BETA WORKSPACES

Workspaces ›
help-gatk/GATK4-Germline-Preprocessing-VariantCalling-JointCalling (read ...

Cloud Environment
None

DASHBOARD | DATA | NOTEBOOKS | WORKFLOWS | JOB HISTORY

ABOUT THE WORKSPACE

#### GATK Best Practices for Germline SNPs & Indels

This workspace contains tutorial notebooks and workflows that cover pre-processing, SNP and Indel variant calling.

There are two ways to run the workflows, in the single sample case one sample is run through the first two workflows and a single VCF file for the sample is generated. In the cohort case, all four workflows are run on several samples to generate a multi-sample VCF. The workflows are properly configured in this workspace to run back to back, so that outputs from each step will automatically become the inputs for the next.

Each workflow follows the GATK Best Practices on human whole-genome sequence data. Detailed description of the workflows is available in Gatk's Best Practices Document.

To learn more about how GATK workflows and Best Practices are used for production at the Broad Institute, you can also view the Whole Genome Analysis Pipeline Workspace.

The workspace material is provided by the GATK team. Please post any questions or concerns to one of our forum sites : GATK or Terra

#### Notebooks

Here you will be running two different tutorial notebooks describing variant calling and different methods for variant filtering. This workspace is read-only, so clone your own unique copy to work with it.

You should find the documentation within the notebooks sufficient to run through them again on your own or to share with a friend later (and we encourage you to do so!).

The notebook(s) in this workspace are:

- 1-germline-variant-discovery-tutorial
- 2-gatk-hard-filtering-tutorial

All notebooks in this workspace can use the following runtime settings:

| Option | Value |
|---|---|
| Environment | Default (GATK 4.1.4.1) |
| Profile | Custom |
| CPU Minimum | 4 |
| Disksize Minimum | 100 GB |
| Memory Minimum | 15 GB |

#### Workflows overview

1. **Preprocessing-For-Variant-Discovery**: takes as input an unmapped BAM list file (text file containing paths to unmapped bam files) to perform preprocessing tasks such as mapping, marking duplicates, and base recalibration. It produces a single BAM file
2. **Haplotypecaller**: takes as input a bam file and produces a file GVCF ( precursor to a VCF )
3. **Generate-Sample-Map**: takes as input several GVCF files and generates a sample map file, which is text file where the first coloumn is the name of the sample and the second column contains the path to the samples GVCF file
4. **Joint-Genotyping**: takes as input a sample map file to perform variants calling all the provided GVCF files and filtering to produce a multi-sample VCF (minimum of 50 samples is required)

Scroll down for details on each workflow, including input and output descriptions and requirements, estimated run times and costs, and information on sample data.

##### Workflow Naming Schema

There are two complete sets of workflows. Those with a "1" in front use hg38 reference data and those that begin with a "2" use b37. The table below lists the workflows and the corresponding reference.

| Workflow Name | Reference Data |
|---|---|
| 1-1-Preprocessing-For-Variant-Discovery | hg38 |
| 1-2-Haplotypecaller | hg38 |
| 1-3-Generate-Sample-Map | hg38 |
| 1-4-Joint-Genotyping | hg38 |
| 2-1-Preprocessing-For-Variant-Discovery | b37 |
| 2-2-Haplotypecaller | b37 |
| 2-3-Generate-Sample-Map | b37 |
| 2-4-Joint-Genotyping | b37 |

##### Input and output data files overview

**Single sample case**
This will produce a VCF file for a single sample.



WORKSPACE INFORMATION

| CREATION DATE | LAST UPDATED |
|---|---|
| 1/13/2020 | 4/1/2021 |
| SUBMISSIONS 4 | ACCESS LEVEL Reader |
| GOOGLE PROJECT ID help-gatk | |

OWNERS

TAGS

best-practices, gatk, germline, HaplotypeCaller, INDEL, Joint Genotyping, pre-processing, SNP, variant calling, variant discovery

Google Bucket
Name: fc-499b7436-38bc-4fb9-b3c1-...
Location: multi-region: US
Open in browser

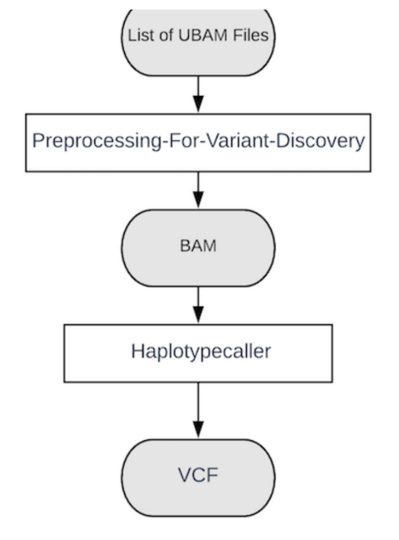


To run the workflows in single sample mode:

1. Head to the workflows tab
2. Click on 1-1-Preprocessing-For-Variant-Discovery-HG38 workflow
   A. Click on "Select Data"
   B. Select "Choose existing sets"
   C. Check the row with "1kgp-50-wgs" as the sample_set
   D. Click "OK"
3. Click on Run analysis
4. Once the workflow has completed head to the Workflows tab and on to the next workflow.
5. Click on 1-2-Haplotypecaller-HG38 workflow
   A. Click on "Select Data"
   B. Select "Choose existing sets"
   C. Check the row with "1kgp-50-wgs" as the sample_set
   D. Click "OK"
   E. In the Input tab set the "make_gvcf" parameter to `False` F. In the output tab set the output_vcf and output_vcf_index parmeter to `this.downsampled_hg38_vcf` and `this.downsampled_hg38_vcf_index`
6. Click on Run analysis

**Cohort sample case**

It's possible to generate a multi-sample VCF instead of a single sample VCF by setting the "make_gvcf" parameter in the **Haplotypecaller** workflow to `True`. Haplotypecaller will then create a GVCF which can be used in the preceding workflows to create the multisample VCF. Running all four workflows takes an unmapped BAM (uBAM) input file and returns a multi-sample VCF. The first two workflows are run once for each sample.

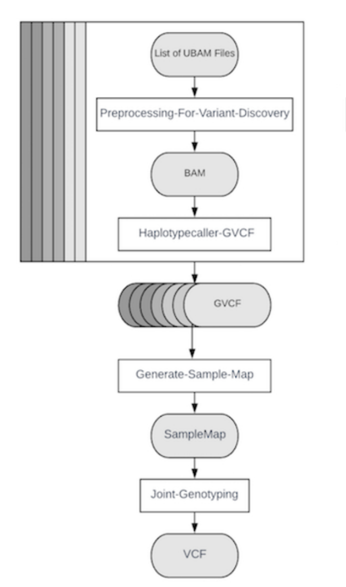


To run the workflows in cohort mode:

1. Head to the workflows tab
2. Click on 1-1-Preprocessing-For-Variant-Discovery-HG38 workflow
   A. Click on "Select Data"
   B. Select "Choose existing sets"
   C. Check the row with "1kgp-50-wgs" as the sample_set
   D. Click "OK"
3. Click on Run analysis
4. Once the workflow has completed head to the Workflows tab and on to the next workflow.
5. Click on 1-2-Haplotypecaller-HG38 workflow

5. Click on 1-2-Haplotypecaller-HG38 workflow
   A. Click on "Select Data"
   B. Select "Choose existing sets"
   C. Check the row with "1kgp-50-wgs" as the sample_set
   D. Click "OK"
   E. Set the "make_gvcf" parameter to `True` F. In the output tab set the output_vcf and output_vcf_index parmeter to `this.downsampled_hg38_gvcf` and `this.downsampled_hg38_gvcf_index`
6. Click on Run analysis
7. Once the workflow has completed head to the Workflows tab and on to the next workflow.
8. Click on 1-3-Generate-Sample-Map-HG38 workflow
   A. Click on "Select Data"
   B. Check the row with "1kgp-50-wgs" as the sample_set
   C. Click "OK"
9. Click on Run analysis
10. Once the workflow has completed head to the Workflows tab and on to the next workflow.
11. Click on 1-4-Joint-Genotyping-HG38 workflow
    A. Click on "Select Data"
    B. Check the row with "1kgp-50-wgs" as the sample_set
    C. Click "OK"
12. Click on Run analysis

If you would like to run the workflows again from scratch, delete the sample and sample_set table and re-upload both tables from this google bucket.

#### Workspace Data

There are two sets of sample tables in the DATA tab - one corresponding to template input files, and another with examples of processed data (i.e. example_sample_output).

**Template input data**
These are the tables you would use to run the workflow pipeline from the beginning.

- participant - Individual per row. This won't be used directly in the workspace but it's best practice to include this in the DATA tab.
- sample - Sample from a participant per row.
  This will be used by the preprocessing and haplotypecaller workflows (*-1 and *-2).
- sample_set - A set of samples per row. This will be used by the Generate-sample-map and joint genotyping workflows (*-3 and *-4)

**Examples of processed data**
Once all the workflows have been executed you will see several additional columns within the sample and sample_set tables. Examples of what you should see are in the following tables:

- example_sample_output
- example_sample_set_output

#### Workflows

##### 1-Preprocessing-For-Variant-Discovery

**What does it do?**
This WDL takes sequencing data in unmapped BAM (uBAM) format and outputs a clean BAM file and its index, suitable for variant discovery analysis.

**What does it require as input?**
The 1-Preprocessing-For-Variant-Discovery workflow accepts a file containing a list of unaligned BAMS. To learn more about how to generate a list file, see this article.

The input data are samples:

- Pair-end sequencing data in unmapped BAM (uBAM) format
- One or more read groups, one per uBAM file, all belonging to a single sample (SM)

Input uBAM files must comply with the following requirements:

- Filenames all have the same suffix (we use ".unmapped.bam")
- Files must pass validation by ValidateSamFile
- Reads are provided in query-sorted order
- All reads must have an RG tag
- Reference index files must be in the same directory as source (e.g. reference.fasta.fai in the same directory as reference.fasta)

**If your sequencing data is not in uBAM format** (e.g. FASTQ), check out this file conversion workspace, https://app.terra.bio/#workspaces/help-gatk/Sequence-Format-Conversion for workflows to convert:

1. Interleaved FASTQ to paired FASTQ
2. Paired FASTQ to unmapped BAM
3. BAM to unmapped BAM
4. CRAM to BAM files from sequencer output for use in GATK analysis tools

**Sample data description and location**
The workspace DATA tab contains downsampled 1000 Genome Project unaligned BAM list files in the `sample` table under the column `flowcell_unmapped_bams_list`.

**What does it return as output?**
The workflow generates a clean BAM file and its index, suitable for variant discovery analyses and stored in the workspace bucket. Metadata for all outputs are written to the `sample` table in the workspace DATA tab.

**Reference data description and location**
Required and optional references and resources for the workflows are included in the Workspace DATA tab in the Reference Data tables. The input unmapped BAM samples have yet to be aligned to a reference so they are not restricted to a particular reference. Only after the first workflow will the samples be restricted to working with a reference, because the unmapped BAM files will be mapped after that point. Again

there are two sets of each workflow, one set configured with hg38 references and the other set configured with b37 references.

**Estimated time and cost to run on sample data**

| Sample Name | Sample Size | Time | Cost $ |
|---|---|---|---|
| NA12878_24RG_small | 3.11 GB | 1:28:00 | 0.18 |
| NA12878 | 64.89 GB | 22:35:00 | 4.98 |
| downsampled-1kgp-50-exomes | 32.13 GB | 02:07:00 | 7.29 |

#### 2-Haplotypecaller

**What does it do?**
The workflow scatters the HaplotypeCaller tool over a sample (clean BAM file and index, from the previous step) using an intervals list file. In particular, it runs the HaplotypeCaller tool from GATK4 on a single sample according to GATK Best Practices. The output file produced will be a single VCF or GVCF file depending on the mode in which it is run. If a GVCF is produced then it can be used by the joint genotyping workflow.

**What does it require as input?**
The workflow accepts: In particular:

- One analysis-ready BAM file for a single sample (as identified in RG:SM), pre-processed using GATK Best Practices.
- A file containing a set of variant calling interval lists for the scatter

**What does it return as output?**
One VCF or GVCF file and its index

**Sample data description and location**
Links to the expected input types are available in the `sample` data table (processed example) for testing. The `sample` data table lists analysis-ready BAM files under the `analysis_ready_bam` column.

**Reference data description and location**
Required and optional references and resources for the workflows are included in the Workspace DATA tab in the Reference Data tables. The **1-2_Haplotypecaller** workflow is configured with HG38 references and **2-2_Haplotypecaller** is configured with B37 references.

**Estimated time and cost to run on sample data**

| Sample Name | Sample Size | Time | Cost $ |
|---|---|---|---|
| NA12878_24RG_small | 4.66 GB | 02:28:00 | 0.21 |
| NA12878 (CRAM) | 19.55 GB | 14:05:00 | 2.24 |
| NA12878 (BAM) | 68.00 GB | 03:44:00 | 1.37 |
| downsampled-1kgp-50-exomes | 23.00 GB | 01:12:00 | 24.25 |

#### 3-Generate-Sample-Map

**What does it do?**
This WDL generates a sample_map file, which can be used for the Joint-Genotyping workflow. A sample map is a tab-delimited text file of 2 columns; 1. the name of the sample and 2. the file path (in this case the Google bucket path of the file).

**What does it require as input?**

- An array of file names
- An array of file paths
- Name of output sample_map

**What does it return as output?**

- Sample map file

**Reference data description and location**
Reference files are not used in this workflow.

#### 4-Joint-Genotyping

**What does it do?**
This WDL implements the joint calling and variant quality score recalibration (VQSR) filtering portion of the GATK Best Practices.

**What does it require as input?**

- GVCFs produced by HaplotypeCaller in GVCF mode
- Bare minimum: 50 samples. Gene panels are not supported

**What does it return as output?**
A VCF file and its index, filtered using VQSR, with genotypes for all samples present in the input VCF. All sites that are present in the input VCF are retained. Filtered sites are annotated as such in the FILTER field.

**Sample data description and location**
Links to the expected input types are available in the processed example data table for testing. The 4-Joint-Genotyping workflow accepts one or more GVCFs produced by haplotypecaller. The `gvcf` column in the data table contains a full-sized GVCF of NA12878 that will be used for the Joint-Genotyping workflow.

**Reference data description and location**
Required and optional references and resources for the workflows are included in the Workspace DATA tab in the Reference Data tables. **1-4_JointGenotyping** is configured with hg38 references and **2-4_JointGenotyping** is configured with the b37 reference.

**Estimated time and cost to run on sample data**

| Sample Name | Sample Size | Time | Cost $ |
|---|---|---|---|
| downsampled-1kgp-50-exomes | 9.52 GB | 03:17:00 | 1.35 |

#### Optional Workflows

Additional workflows have been added for your convenience.

**Optional-Paired-FASTQ-to-Unmapped-BAM**: This WDL converts paired FASTQ to uBAM and adds read group information.

Requirements/expectations

- Pair-end sequencing data in FASTQ format (one file per orientation)
- The following metadata descriptors per sample:
  - readgroup
  - sample_name
  - library_name
  - platform_unit
  - run_date
  - platform_name
  - sequecing_center Outputs
  - Unmapped BAM

**Optional-Gatk-GatherVCFsCloud**: This tool combines together rows of variant calls from multiple VCFs, e.g. those produced by scattering calling across genomic intervals, into a single VCF. This tool enables scattering operations, e.g. in the cloud, and is preferred for such contexts over Picard MergeVcfs or Picard GatherVCfs. The input files need to have the same set of samples but completely different sets of loci. These input files must be supplied in genomic order and must not have events at overlapping positions.

Input

- A set (array) of VCF files, sorted by genomic position. Its also possible to provide a file containing a list of VCF files, each row being an individual VCF file path.

Output

- A single VCF file containing the variant call records from the multiple VCFs.

**Optional-ReblockGVCF-gatk4_exomes_goodCompression**: Users working with large sample sets can invoke the GnarlyGenotyper task in the JointGenotyping.wdl workflow. However, the ReblockGVCF tool must be run for all GVCFs produced by HaplotypeCaller before they can be appropriately processed by GnarlyGenotyper.

Input

- A GVCF file

Output

- Reblocked GVCF file

#### Important notes on workflow limitations

**Small Cohorts**

We believe the results of this workflow run on a single WGS sample are equally accurate, but there may be some shortcomings when the workflow is modified and run on small cohorts. Specifically, modifying the SNP ApplyRecalibration step for higher specificity may not be effective. You can verify if this is an issue by consulting the gathered SNP tranches file. If the listed `truthSensitivity` in the rightmost column is not well matched to the `targetTruthSensitivity` in the leftmost column, then requesting that `targetTruthSensitivity` from ApplyVQSR will not use an accurate filtering threshold.

Additionally

- No allele subsetting for the Joint-Genotyping workflow
  - For large cohorts, even exome callsets can have more than 1000 alleles at low complexity/STR sites
  - For sites with more than six alternate alleles (by default) called genotypes will be returned, but without the PLs since the PL arrays get enormous
  - Allele-specific filtering could be performed if AS annotations are present, but the data will still be in the VCF in one giant INFO field
- JointGenotyping output is divided into lots of shards
  - Desirable for use in Hail, which supports parallel import
  - It's possible to use GatherVcfs to combine shards.
- GnarlyGenotyper uses a QUAL score approximation
  - Dramatically improves performance compared with GenotypeGVCFs, but QUAL output (and thus the QD annotation) may be slightly discordant between the two tools.

**Exomes**

Currently the workflows are configured for WGS processing. The dynamic scatter interval creating task was optimized for genomes. The scattered SNP VariantRecalibration may fail because of too few "bad" variants to build the negative model. Also, apologies that the logging for SNP recalibration is overly verbose.

The provided tool configurations are meant to be a ready-to-use example of the workflows. It is the user's responsibility to correctly set the reference and resource input variables using the GATK Tool and Tutorial Documentations.

**GnarlyGenotyper**

Users working with large sample sets can invoke the GnarlyGenotyper task in the JointGenotyping.wdl workflow. However, the ReblockGVCF tool must be run for all GVCFs produced by HaplotypeCaller before they can be appropriately processed by GnarlyGenotyper. A workflow that applies the reblocking tool is provided here: ReblockGVCF-gatk4_exomes_goodCompression

**Controlling cloud costs**

Note that cost and time estimates will vary with the use of preemptibles. Using preemptibles can save up to 80% on compute costs. For further helpful hints on controlling cloud costs, see this article and for additional ways to estimate cloud cost use Google's cost calculator.

#### Software Versions

- GATK 4.1.4.0

- BWA 0.7.15-r1140
- Picard 2.16.0-SNAPSHOT
- Samtools 1.3.1 (using htslib 1.3.1)
- Python 2.7

---

##### Contact information

For questions about this workspace please visit the Featured Workspaces Topic topic on the [Terra Community Forum](). Use the search box to see if other users have asked the same question previously. If not, post and tag **@Beri Shifaw** so that we get notified.

This material is provided by the GATK Team. Please post any questions or concerns regarding the GATK tool to the [GATK forum]()

##### License

**Copyright Broad Institute, 2019 | BSD-3**
All code provided in this workspace is released under the WDL open source code license (BSD-3) (full license text at https://github.com/openwdl/wdl/blob/master/LICENSE). Note however that the programs called by the scripts may be subject to different licenses. Users are responsible for checking that they are authorized to run all programs before running these tools.

##### Workspace Change Log

| Date | Change | Author |
|---|---|---|
| 2020-01-14 | Initial feature of workspace | Beri Shifaw |
| 2020-02-24 | Added Optional-ReblockGVCF-gatk4_exomes_goodCompression workflow | Beri Shifaw |
| 2020-04-02 | Updated Preprocessing-For-Variant-Discovery to v2.0.0, Reuploaded HC and Generate Map workflow | Beri Shifaw |
| 2020-06-22 | Added workflow overview image to dashboard and additional description to workflows, Updated Joint Genotyping workflow, removed "GVCF" from Haplotypecaller name to indicate the workflow can be run to create VCF | Beri Shifaw |
| 2020-09-28 | Updated Haplotypecaller and JointGenotype workflow to release 2.2.0 | Beri Shifaw |
| 2020-11-28 | Updated Haplotypecaller to release 2.3.0, updated JointGenotype workflow source to Warp repo, added GATK Notebook tutorials | Beri Shifaw |
| 2020-12-07 | Updated data tables to use downsample wgs | Beri Shifaw |
| 2021-01-29 | Updated dashboard to refer to [Whole Genome Analysis Pipeline Workspace]() | Beri Shifaw |

###### DATASET ATTRIBUTES

---

Structured Data Use Limitations

Supplemental Note 2. Analyzing differential gene expression with Bioconductor's edgeR

POWERED BY Terra BETA WORKSPACES

Workspaces ›
help-gatk/Bioconductor (read only)

Cloud Environment
None

DASHBOARD DATA NOTEBOOKS WORKFLOWS JOB HISTORY

##### ABOUT THE WORKSPACE

Explore common Bioconductor packages that can be used to perform bulk RNA differential expression analyses or manipulate single-cell RNA-seq data

### Table of Contents


- Getting started with Bioconductor in Terra
- Using the Bioconductor workspace
  - Cloud Environment configuration and cost
  - Testing for differential gene expression using the "edgeR" Notebook
    - Sample data
    - Required tools (R packages)
    - Analyses
  - Exploring single-cell RNAseq data using the 'SingleCellExperiment' Notebook
    - Sample data
    - Required tools (R packages)
    - Analyses


### Getting started with Bioconductor in Terra

Bioconductor is a suite of open source tools, primarily written as R packages, designed for the statistical analysis of high-throughput genomic data. The Terra Bioconductor image is an extension of the Terra R image and comes preloaded with commonly used Bioconductor packages. These include tools for:

1. Analyzing single cell sequencing data (SingleCellExperiment)
2. Annotating genes (GenomicFeatures)
3. Storing and manipulating short genomic alignments (GenomicAlignments)
4. Manipulating and assessing the quality of FASTQ files (ShortRead)
5. Performing RNA-seq differential gene expression analyses (DESeq2).

Don't see the package you need? No problem! You can download ANY Bioconductor package using the `BiocManager::install()` command. View additional pre-installed Bioconductor packages and Docker image details in the Terra Bioconductor ReadMe.

You can also view all the contents of the Bioconductor image by selecting the Cloud Environment and choosing "What's installed on this environment?" (shown below). To identify Bioconductor-relevant packages, choose R in the Installed packages dropdown.

| Viewing Bioconductor packages | Image |
|---|---|
| Select "What's installed on this environment" | 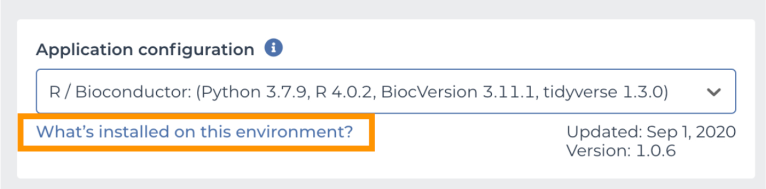 |
| Select "R " in dropdown | 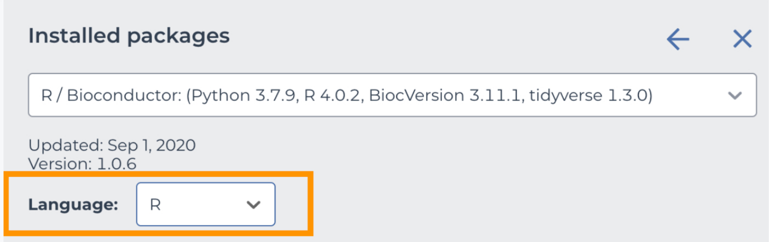 |

### Using the Bioconductor workspace

##### WORKSPACE INFORMATION

| CREATION DATE | LAST UPDATED |
|---|---|
| 4/22/2020 | 4/2/2021 |

| SUBMISSIONS | ACCESS LEVEL |
|---|---|
| 0 | Reader |

GOOGLE PROJECT ID
help-gatk

##### OWNERS

##### TAGS

Bioconductor edgeR RNAseq single-cell

Google Bucket
Name: fc-0fe0fb8e-70f6-412b-8f3c-0...
Location: multi-region: US
Open in browser

This workspace uses Jupyter Notebooks to showcase two commonly used Bioconductor packages designed for RNAseq analyses: 'edgeR' and 'SingleCellExperiment'.

**To get started with these notebooks, make a clone of this workspace** using the three button icon in the upper right of the dashboard. Then read through the following sections to learn more about the workspace configurations, Notebooks, example data and analyses.

#### Cloud environment configuration and cost

To use the preset Bioconductor image in a new workspace, first **click on the Cloud Environment icon (top right of the new workspace's dashboard)** and set the environment and compute power as follows:

<table>
<tr><th>Environment</th><th>Screenshot</th></tr>
<tr><td>Select the Bioconductor option from the Environment drop-down.</td><td>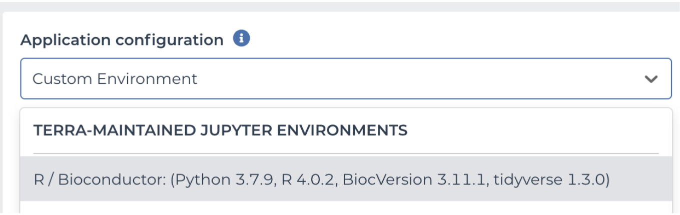
</td></tr>
</table>

<table>
<tr><th>Compute power</th><th>Screenshot</th></tr>
<tr><td>Use the default standard VM settings, which will cost approximately $0.19 per hour.</td><td>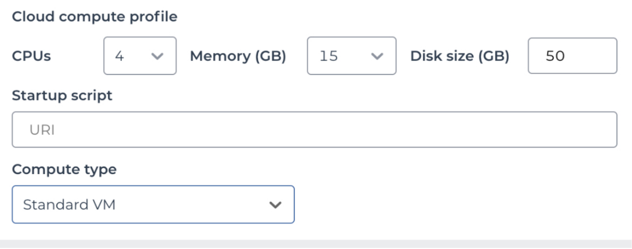
</td></tr>
</table>

---

#### Testing for differential gene expression using the "edgeR" Notebook

The "edgeR" Notebook uses the Bioconductor edgeR package to analyze publically available bulk RNAseq gene count data.

##### Sample data

The sample data used for this Notebook are read counts derived from [Sato et al. 2020] ([GEO #GSE124407]). The counts are uploaded to a public Google Bucket as a tab-delimited text file ("counts.txt") that contains a gene ID column followed read count columns for six samples. Instructions and sample code for accessing Google Bucket files from a Jupyter Notebook are listed in this edgeR Notebook.

##### Required tools (R packages)

This Notebook uses the following R packages:

- 'methods': common methods and classes for R objects
- 'edgeR': statistical tools for RNAseq differential expression analyses
- 'RColorBrewer': color schemes for graphics
- 'ggplot2': a data visualization package

##### Analyses

This Notebook performs the following:

- Creates an edgeR DGEList object
- Filters and normalizes count data
- Estimates a common negative binomial dispersion parameter
- Performs a likelihood ratio test (LRT) for differential expression
- Creates smear plot of differential expression

#### Exploring single-cell RNAseq data using the 'SingleCellExperiment' Notebook

The "SingleCellExperiment" Notebook uses Bioconductor's SingleCellExperiment package, which defines a container for storing single-cell RNAseq data. This includes common gene and cell metadata, such as size factors, control RNA transcript spike-ins, and antibody/CRISPR tags.

##### Sample data

The sample data for this Notebook is the Allen Brain Single-cell RNAseq subset stored in the Bioconductor package 'scRNAseq'. Details about using this dataset can be found in the Notebook.

##### Required tools (R packages)

This Notebook uses the following pre-installed R packages:

- 'SingleCellExperiment': S4 class for storing single-cell data
- 'scRNAseq': package of gene-level counts for publically available RNAseq datasets
- 'Rtsne': R wrapper around the T-distributed Stochastic Neighbor Embedding implementation
- 'magrittr': package that provides a handy pipe-like function
- 'ggplot2': a data visualization package

##### Analyses

This Notebook performs the following:

- Creates a SingleCellExperiment object
- Labels spike-in transcripts
- Accounts for sequencing depth
- Makes a SingleCellExperiment subset
- Plots a SingleCellExperiment subset in a t-SNE

---

---

#### Additional Resources

You can find additional resources for using Terra in the Terra Support and read more about the Bioconductor Image in "Using the Bioconductor Docker image in Terra".

#### License

**Copyright Broad Institute, 2019 | BSD-3**
All code provided in this workspace is released under the WDL open source code license (BSD-3) (full license text at https://github.com/openwdl/wdl/blob/master/LICENSE). Note however that the programs called by the scripts may be subject to different licenses. Users are responsible for checking that they are authorized to run all programs before running these tools.

#### Questions and Feedback

Please post workspace questions and feedback to the Featured Workspaces community forum (login required). Tag @Liz Kiernan and @Anton Kovalsky in the “Details” section of your post.

### Analyzing Differential Gene Expression with Bioconductor's 'edgeR'

#### Overview

This tutorial explores how to quantify differential gene expression using Bioconductor's 'edgeR' package. It uses a modified read counts file from publically available dataset generated from human pancreatic ductal adenocarcinoma cells (AsPC-1) lacking BACH1, a transcription factor invovled in oxidative stress pathways (Sato et al. 2020; GEO #GSE124407).

We will use this tutorial's example dataset to examine differential gene expression between control and BACH1 knockout cells.

### Set-up

#### Load packages

First, we will load the necessary packages using the library() function.

***Note about warnings*** You may get a pink warning after running a code block. These warnings are usually just that - warnings - and do not break the analysis.

In [1]:
```
# Load the necessary libraries
library("methods")
library("edgeR")
library("RColorBrewer")
library("ggplot2")
```

```
Loading required package: limma
```

#### Access gene count data from a Google bucket

The example read counts are listed in a tab-delimited text file ("counts.txt") located in a public Google bucket. However, if you would like to try these analyses on your own data, you can upload your count data to the workspace bucket. The first column of the count file should contain a gene ID list and the subsequent columns should contain the read counts for each sample.

**To use the example (public-access) counts file**

Copy the file (which is stored in a public Google bucket) to your virtual machine using the "system(paste0())" command provided in the code block below.

**To use your own counts file**

Upload your own data to your workspace bucket by going to the workspace Data tab, selecting Files, and clicking the upload icon.

To access these files from this Notebook, we use the gsutil command. For this command, you need to specify the Google bucket location (IRL starting with "gs://..."). If you are using your own workspace bucket instead of the public bucket, you can find the Google bucket id on the right side of the workspace dashboard.

In [2]:
```
# Access counts file in public bucket
system(paste0("gsutil cp gs://terra-featured-workspaces/Bioconductor/*.txt ."))

# Access counts file in your workspace bucket by uncommenting the command "system(paste())" below
# Note that you need to change to your workspace bucket path...
# system(paste0("gsutil cp gs://your_workspace_bucket/*.txt ."))
```

In [3]:
```
# Sanity check - list all files in the notebook directory to verify that the counts.txt file is there
list.files()
```

'counts.txt' · 'edgeR.ipynb' · 'SingleCellExperiment.ipynb'

#### Read the count file

Let's examine the contents of the text file to ensure it contains our count data. To do this, use the read.table() function. After running the following command, notice the text file has 7 columns. The first is the gene_id (Ensembl IDs), the next three are read counts for each control sample ("sgC"), and the last three are gene counts for each BACH1 knockout samples ("sg2").

In [4]:
```
read.table('counts.txt', header = TRUE, sep = "", dec = ".",stringsAsFactors=F)
```

A data.frame: 57820 × 7

| id | sgC.1 | sgC.2 | sgC.3 | sg2.1 | sg2.2 | sg2.3 |
|---|---|---|---|---|---|---|
| <chr> | <int> | <int> | <int> | <int> | <int> | <int> |
| ENSG00000000003.10 | 513 | 342 | 459 | 969 | 1046 | 1390 |
| ENSG00000000005.5 | 0 | 0 | 0 | 0 | 0 | 0 |
| ENSG00000000419.8 | 600 | 464 | 517 | 735 | 795 | 1035 |
| ENSG00000000457.9 | 245 | 222 | 164 | 293 | 355 | 285 |
| ENSG00000000460.12 | 520 | 398 | 388 | 504 | 437 | 507 |

| | | | | | | |
|---|---|---|---|---|---|---|
| [illegible] | [illegible] | [illegible] | [illegible] | [illegible] | [illegible] | [illegible] |
| ENSG00000000938.8 | 0 | 2 | 1 | 0 | 0 | 2 |
| ENSG00000000971.11 | 373 | 185 | 209 | 5 | 0 | 3 |
| ENSG00000001036.9 | 1169 | 870 | 1067 | 1006 | 973 | 1230 |
| ENSG00000001084.6 | 1319 | 1044 | 1061 | 2823 | 2571 | 3259 |
| ENSG00000001167.10 | 1062 | 884 | 1036 | 855 | 831 | 1118 |
| ENSG00000001460.13 | 294 | 256 | 247 | 232 | 248 | 291 |
| ENSG00000001461.12 | 1581 | 999 | 1204 | 1196 | 1095 | 1516 |
| ENSG00000001497.12 | 828 | 718 | 699 | 1173 | 1157 | 1726 |
| ENSG00000001561.6 | 65 | 50 | 4 | 49 | 45 | 51 |
| ENSG00000001617.7 | 15 | 8 | 22 | 15 | 9 | 13 |
| ENSG00000001626.10 | 0 | 0 | 0 | 2 | 1 | 2 |
| ENSG00000001629.5 | 3234 | 2147 | 2548 | 2272 | 2267 | 2795 |
| ENSG00000001630.11 | 241 | 228 | 289 | 319 | 326 | 343 |
| ENSG00000001631.10 | 736 | 699 | 608 | 645 | 658 | 749 |
| ENSG00000002016.12 | 249 | 124 | 204 | 188 | 204 | 154 |
| ENSG00000002079.8 | 72 | 161 | 135 | 56 | 31 | 100 |
| ENSG00000002330.9 | 832 | 511 | 699 | 705 | 652 | 753 |
| ENSG00000002549.8 | 1042 | 590 | 771 | 750 | 895 | 970 |
| ENSG00000002586.13 | 578 | 368 | 418 | 576 | 518 | 768 |
| ENSG00000002587.5 | 274 | 241 | 243 | 284 | 306 | 478 |
| ENSG00000002726.15 | 262 | 141 | 206 | 367 | 357 | 471 |
| ENSG00000002745.8 | 130 | 54 | 87 | 19 | 16 | 27 |
| ENSG00000002746.10 | 119 | 48 | 122 | 32 | 52 | 52 |
| ENSG00000002822.11 | 630 | 491 | 706 | 540 | 573 | 706 |
| ENSG00000002834.13 | 12112 | 8755 | 11130 | 8564 | 8543 | 10735 |
| ⋮ | ⋮ | ⋮ | ⋮ | ⋮ | ⋮ | ⋮ |
| ENSGR0000198223.10 | 0 | 0 | 0 | 0 | 0 | 0 |
| ENSGR0000205755.6 | 0 | 0 | 0 | 0 | 0 | 0 |
| ENSGR0000214717.5 | 0 | 0 | 0 | 0 | 0 | 0 |
| ENSGR0000223274.1 | 0 | 0 | 0 | 0 | 0 | 0 |
| ENSGR0000223484.2 | 0 | 0 | 0 | 0 | 0 | 0 |
| ENSGR0000223511.1 | 0 | 0 | 0 | 0 | 0 | 0 |
| ENSGR0000223571.1 | 0 | 0 | 0 | 0 | 0 | 0 |
| ENSGR0000223773.2 | 0 | 0 | 0 | 0 | 0 | 0 |
| ENSGR0000225661.2 | 0 | 0 | 0 | 0 | 0 | 0 |
| ENSGR0000226179.1 | 0 | 0 | 0 | 0 | 0 | 0 |
| ENSGR0000227159.3 | 0 | 0 | 0 | 0 | 0 | 0 |
| ENSGR0000228410.1 | 0 | 0 | 0 | 0 | 0 | 0 |
| ENSGR0000228572.2 | 0 | 0 | 0 | 0 | 0 | 0 |
| ENSGR0000229232.1 | 0 | 0 | 0 | 0 | 0 | 0 |
| ENSGR0000230542.1 | 0 | 0 | 0 | 0 | 0 | 0 |
| ENSGR0000234622.1 | 0 | 0 | 0 | 0 | 0 | 0 |
| ENSGR0000234958.1 | 0 | 0 | 0 | 0 | 0 | 0 |
| ENSGR0000236017.3 | 0 | 0 | 0 | 0 | 0 | 0 |
| ENSGR0000236871.2 | 0 | 0 | 0 | 0 | 0 | 0 |
| ENSGR0000237040.1 | 0 | 0 | 0 | 0 | 0 | 0 |
| ENSGR0000237531.1 | 0 | 0 | 0 | 0 | 0 | 0 |
| ENSGR0000237801.1 | 0 | 0 | 0 | 0 | 0 | 0 |
| ENSGR0000263835.1 | 0 | 0 | 0 | 0 | 0 | 0 |
| ENSGR0000263980.1 | 0 | 0 | 0 | 0 | 0 | 0 |
| ENSGR0000264510.1 | 0 | 0 | 0 | 0 | 0 | 0 |
| ENSGR0000264819.1 | 0 | 0 | 0 | 0 | 0 | 0 |
| ENSGR0000265350.1 | 0 | 0 | 0 | 0 | 0 | 0 |
| ENSGR0000265658.1 | 0 | 0 | 0 | 0 | 0 | 0 |
| ENSGR0000266731.1 | 0 | 0 | 0 | 0 | 0 | 0 |
| ENSGR0000270726.1 | 0 | 0 | 0 | 0 | 0 | 0 |

#### Prepare a count dataframe

We can set up a few variables for downstream analyses. In addition to specifying the sample and column names, we will name the count dataframe as the variable 'all.data'.

In [5]:
```
# Specify sample names and column labels in the dataframe
samples <- c("sgC-1","sgC-2","sgC-3","sg2-1","sg2-2","sg2-3")
all.data <- read.delim("counts.txt", col.names=c("gene_id", "sgC-1","sgC-2","sgC-3","sg2-1","sg2-2","sg2-3"),  sep
="\t")
colnames(all.data)[2:ncol(all.data)] <- samples
```

```
# Sanity check - View the top lines
head(all.data)
```

A data.frame: 6 × 7

| | gene_id | sgC-1 | sgC-2 | sgC-3 | sg2-1 | sg2-2 | sg2-3 |
|---|---|---|---|---|---|---|---|
| | <fct> | <int> | <int> | <int> | <int> | <int> | <int> |
| **1** | ENSG00000000003.10 | 513 | 342 | 459 | 969 | 1046 | 1390 |
| **2** | ENSG00000000005.5 | 0 | 0 | 0 | 0 | 0 | 0 |
| **3** | ENSG00000000419.8 | 600 | 464 | 517 | 735 | 795 | 1035 |
| **4** | ENSG00000000457.9 | 245 | 222 | 164 | 293 | 355 | 285 |
| **5** | ENSG00000000460.12 | 520 | 398 | 388 | 504 | 437 | 507 |
| **6** | ENSG00000000938.8 | 0 | 2 | 1 | 0 | 0 | 2 |

#### Prepare a count dataframe using sample subsets

In this notebook, we are analyzing all of the samples listed in our counts file, but if we wanted to only use a subset of the data, we could specify that subset using the commands below.

After defining the subset, we specify the experimental group to which each sample belongs. In this case, we use "C" for control and "T" for treatment.

We then create a new dataframe (groupDF) containing only the subsetted data.

In [6]:
```
# Make sample groups
sampleGroup=c("sgC-1","sgC-2","sgC-3","sg2-1","sg2-2","sg2-3")

subset<- samples[samples %in% c("sgC-1","sgC-2","sgC-3","sg2-1","sg2-2","sg2-3")]
subset

#Specify the treatment conditions as 'group'
group<-c("C","C","C","T","T","T")
```

'sgC-1' · 'sgC-2' · 'sgC-3' · 'sg2-1' · 'sg2-2' · 'sg2-3'

In [7]:
```
# Reprocess original countdata using the subset
groupDF<-(all.data[, subset])
```

### Create a DGEList Object

The following commands will set up a list-based object called a DGEList using the groupDF dataframe we created. Then, we filter the count data so that our analyses only include genes with a cpm > 1 in at least 3 samples. We normalize our data using the trimmed mean of M-values (TMM) technique. Finally, we visualize the normalized counts in a boxplot.

In [8]:
```
# Use the DGEList function to set up the count dataframe as a DGEList object
yGrp<-DGEList(counts=groupDF, group=group,  genes=all.data[,1])
#Filter the data so that at least 3 treatment groups must have a cpm > 1.
keepGrp<- rowSums(cpm(yGrp)>1) >=3; yGrp <-yGrp[keepGrp, ]
#Normalize the data using the TMM method
yGrp <- calcNormFactors(yGrp, method = "TMM")
#Prepare the data for plotting and make a boxplot
lcpmGrp<-cpm(yGrp, log = TRUE)
boxplot(lcpmGrp)
```

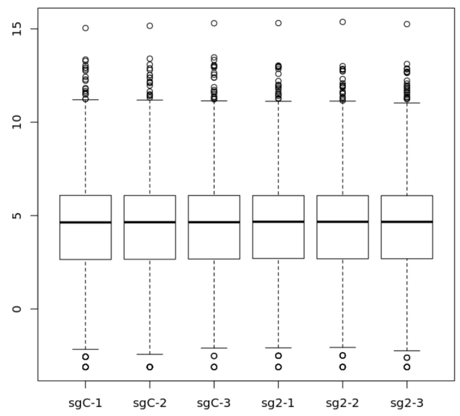


### Create a Design Matrix for the Experimental Conditions

The following commands create a matrix in which each sample is assigned an experimental treatment

In [9]:
```
# Design matrix
design<-model.matrix(~0+yGrp$samples$group, data=yGrp$samples)
colnames(design)<- levels(yGrp$samples$group)
print(design)
```

```
      C T
sgC-1 1 0
sgC-2 1 0
sgC-3 1 0
sg2-1 0 1
sg2-2 0 1
sg2-3 0 1
attr(,"assign")
[1] 1 1
attr(,"contrasts")
attr(,"contrasts")$`yGrp$samples$group`
[1] "contr.treatment"
```

You should now see a matrix in which each sample is assigned to its respective control (C) or treatment (T) group.

### Estimate Dispersion and Test for Differential Expression

We can estimate common, trended, and tag-wise dispersions and then plot each dispersion. We then fit a negative binomial GLM for each tag and perform a likelihood ratio test (LRT) for differential expression between our control samples and BACH1 knockout samples. For this example, we look for differential expression at a false discovery rate (FDR) of 5%.

In [10]:
```
# Esitmate common disp
yGrp<- estimateGLMCommonDisp(yGrp, design, verbose=TRUE)
```

```
Disp = 0.0224 , BCV = 0.1497
```

In [11]:
```
#Estimate trended disp
yGrp <- estimateGLMTrendedDisp(yGrp, design)

#Estimate tag dispersion
yGrp <- estimateGLMTagwiseDisp(yGrp, design)

#Make a plot of the estimated dispersion data
plotBCV(yGrp)
```

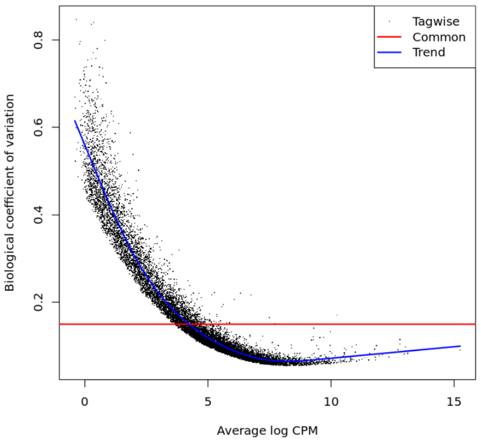


You should see a graph of Average log CPM v. Biological coefficient of variation with tagwise, Common, Trend dispersion represented.

In [12]:
```
# Fit negative binomial GLM for each tag
fit<-glmFit(yGrp, design)
```

pathways (Sato et al. 2020; GEO #GSE124407).

We will use this tutorial's example dataset to examine differential gene expression between control and BACH1 knockout cells.

### Set-up

#### Load packages

First, we will load the necessary packages using the library() function.

***Note about warnings*** You may get a pink warning after running a code block. These warnings are usually just that - warnings - and do not break the analysis.

In [1]:
```
# Load the necessary libraries
library("methods")
library("edgeR")
library("RColorBrewer")
[illegible]
```

13347 · 1

A TestResults:
6 × 1 of type
int

**-1*C 1*T**

| | |
|---|---|
| **1** | 1 |
| **3** | 1 |
| **4** | 0 |
| **5** | 0 |
| **7** | -1 |
| **8** | 0 |

```
       -1*C 1*T
Down        1933
NotSig      9620
Up          1794
```

You should see a display of the number of genes downregulated, unchanged (NotSig) or upregulated in the Treatment group (T)

In [15]:
```
# Direction of differential expression and setting false discovery rate
de<-decideTestsDGE(lrt, p.value = 0.05, adjust.method = "fdr")
summary(de)
```

```
       -1*C 1*T
Down        1933
NotSig      9620
Up          1794
```

If we examine the output above, we see the treatment (BACH1 knockout) decreased the expression of 1,933 genes and increased the expression of 1,794 genes at an FDR of 5%. We can examine the most significant genes by using the topTags(() command.

In [16]:
```
topTags(lrt)
```

**$table**

A data.frame: 10 × 6

| | genes | logFC | logCPM | LR | PValue | FDR |
|---|---|---|---|---|---|---|
| | <fct> | <dbl> | <dbl> | <dbl> | <dbl> | <dbl> |
| **2879** | ENSG00000103888.11 | 6.015327 | 6.595132 | 2210.0375 | 0.000000e+00 | 0.000000e+00 |
| **14183** | ENSG00000176153.10 | 5.706466 | 6.685978 | 1664.8550 | 0.000000e+00 | 0.000000e+00 |
| **2266** | ENSG00000100292.12 | 5.702056 | 7.914835 | 1887.6280 | 0.000000e+00 | 0.000000e+00 |
| **1932** | ENSG00000091129.15 | 4.209553 | 7.133577 | 2066.1231 | 0.000000e+00 | 0.000000e+00 |
| **1892** | ENSG00000090382.2 | 3.623257 | 8.628066 | 1067.2897 | 4.249966e-234 | 1.134486e-230 |
| **4802** | ENSG00000117983.13 | 5.742994 | 5.306808 | 1015.6621 | 7.078120e-223 | 1.574528e-219 |
| **1714** | ENSG00000086548.8 | 2.517185 | 9.888734 | 882.8345 | 5.289849e-194 | 1.008623e-190 |
| **9088** | ENSG00000147689.12 | -2.257501 | 7.889795 | 805.5806 | 3.301671e-177 | 5.508426e-174 |
| **12310** | ENSG00000167767.9 | -3.347514 | 7.289286 | 750.1255 | 3.767946e-165 | 5.587864e-162 |
| **2842** | ENSG00000103485.13 | -3.503298 | 5.818039 | 746.5295 | 2.280439e-164 | 3.043702e-161 |

**$adjust.method**
'BH'
**$comparison**
'-1*C 1*T'
**$test**
'glm'

### Plot Differential Expression

Lastly, we can plot our differential expression using a smear plot.

In [17]:
```
plotMD(lrt)
```

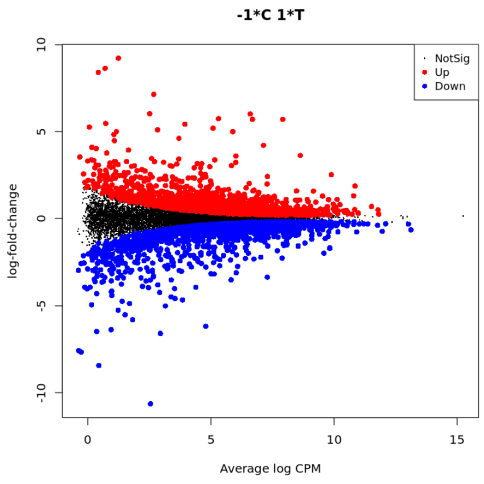


You should see a smear plot in which non-signficant changes are plotted in black, genes upregulated by the treatment (T) relative to control (C) are shown in red, and genes downregulated by the treatment are shown in blue.

This concludes the edgeR tutorial. For more information on edgeR, please see Bioconductor's edgeR page.
